# Physiological signatures but no evidence for universal genomic predictors of extreme drought responses in European beech

**DOI:** 10.64898/2026.09.20.752377

**Authors:** Cheng Li, Lucienne de Witte, Ewa A. Czyż, Aboubakr Moradi, Sabine Braun, Sven-Eric Hopf, Bernhard Schmid, Michael E. Schaepman, Meredith C. Schuman

## Abstract

Following a severe 2018 drought affecting much of Europe, European beech (*Fagus sylvatica*) exhibited patchy canopy dieback. To investigate the drivers of this mortality, we used a paired case-control design across Swiss forests, integrating phenotypic assessments and genomic analyses. Phenotypically, crown transparency and discoloration were used to classify tree damage state, while several other characteristics were measured to evaluate whether paired, neighboring damaged and undamaged trees were otherwise comparable, and to identify physiological stress signatures. Elevated leaf calcium content and red edge reflectance in dried leaves emerged as robust indicators of physiological stress. Genomic association models using reference-based SNP markers and reference-free *k*-mers were heavily confounded by micro-environmental variance and local population stratification even within our geographically limited study region. Our cross-validated genomic outliers showed no spatial overlap with predictive drought-associated loci previously identified in an independent German cohort. These findings indicate that drought survival in European beech is not governed by a universal set of major-effect alleles. Instead, tree performance appears highly context-dependent, likely shaped by micro-environmental heterogeneity and phenotypic condition. However, our results do not exclude polygenic drought responses involving multiple small-effect loci.

## Introduction

Forest ecosystems harbor substantial biodiversity and perform functions critical for human society, from carbon sequestration to soil stabilization. These ecosystems face increasing stress due to a rapidly changing climate. Severe droughts in 2018 caused substantial tree mortality in many common and widespread species throughout central Europe. This trend has since persisted across several years, demonstrating the sustained impact of extreme climate events on forest dynamics (***Schuldt et al., 2020; Braun et al., 2020b; Buras et al., 2020; Rieder et al., 2026***). During these drought events, many trees displayed pronounced signs of stress. Documented symptoms included early leaf discoloration, premature leaf shedding, branch dieback (***Schuldt et al., 2020***), and early fruit abortion (***Nussbaumer et al., 2020***). Low foliar water potential and subsequent xylem hydraulic failure have been identified as primary physiological mechanisms driving damage in several temperate forest tree species under severe soil desiccation (***Dietrich et al., 2019; Vitasse et al., 2019; Braun et al., 2019, 2015; Choat et al., 2018***).

The European beech (*Fagus sylvatica* L.), a keystone forest species across Europe with high economic and societal value, is a common subject of tree ecophysiological and population genetic research (***Magri, 2008; Stefanini et al., 2023***). Genome-environment association studies indicate that gene flow in European beech promotes the spread of advantageous alleles, supporting local populations in adapting to different environmental niches (***Pluess et al., 2016***). Like other widespread temperate forest trees, European beech has maintained large effective population sizes over millennia, providing a reservoir of standing genetic variation that may serve as raw material for adaptation to future climate extremes (***Milesi et al., 2024***). Recently, genome-wide association studies (GWAS) have sought to identify specific genetic variants underlying drought resilience. For instance, Pfenninger and colleagues (***Pfenninger et al., 2021, 2024***) demonstrated that a defined set of single nucleotide polymorphisms (SNPs) could predict drought phenotype (apparently damaged, or apparently healthy) with high accuracy in a German beech cohort. However, European beech is characterized by high phenotypic plasticity (***Gárate-Escamilla et al., 2019***) and shows a geographic gradient in population structure (***Lazic et al., 2024***). It remains unresolved whether the genetic architecture of drought resistance relies on universal, major-effect alleles, or whether it is more localized and context-dependent. Furthermore, it is also possible that drought resistance is highly polygenic and arises from the contribution of many loci with minor-effect alleles whose contributions vary among populations and environments (***Berg and Coop, 2014***), making consistent genome-wide signals difficult to detect in natural cohorts.

At the stand level, small-scale variations in environmental variables, such as soil structure and hydrology, can strongly affect water availability (***Bolte and Villanueva, 2006***). Recent studies in Switzerland indicate that crown damage in *F. sylvatica* increased with decreasing overall water availability, and that more damaged trees often showed lower pre-drought growth or vigor, suggesting that pre-existing tree condition modulates drought vulnerability (***Klesse et al., 2022; Neycken et al., 2022; Braun et al., 2025***). Yet, a prominent observation during recent European droughts is the patchy nature of canopy dieback: severely damaged trees have been frequently found immediately adjacent to apparently healthy trees, occupying the same habitat and of similar age (***Pfenninger et al., 2021; Braun et al., 2021; Klesse et al., 2022; Frei et al., 2022; van der Maaten et al., 2024***). This highly localized mosaic of damage provides a natural experimental framework to test if differences between pairs of undamaged and drought-damaged trees within habitats differ in genotype and micro-environment.

In this study, we employed a paired case-control design in monitored Swiss beech forest stands. We first classified neighboring trees as damaged or healthy based on field assessments of crown transparency and discoloration following the 2018/2019 drought events. We then characterized additional phenotypic and genomic associations with apparent tree health status, combining detailed *in situ* phenotypic assessments—including leaf spectroscopy and foliar nutrient analysis—with dual-tiered genomic sequencing (low-coverage individual sequencing and pooled sequencing). To mitigate reference bias and account for highly structured local populations, we applied both reference-based (SNP) and reference-free (*k*-mer) GWAS testing. By directly comparing our cross-validated genomic outliers with previously published markers **(*Pfenninger et al., 2021, 2024*)**, we aimed to determine whether drought survival in European beech is characterized by universal genetic predictors, more localized genetic structure, environmental sorting, or polygenic drought responses involving multiple small-effect loci.

## Results

### Phenotypic differences between healthy and damaged trees

A phenological analysis of the trees assessed in 2019 has been published by (***Braun et al., 2021***). In this study, we examined each feature separately for the pairs. This analysis aimed to evaluate the experimental pairing of the study trees, assessing whether they differed primarily in their apparent damage status (which was our aim), while acknowledging that recent drought could also affect other phenotypic features. Tree damage status was assigned from field assessments of crown transparency and discoloration in 2019; therefore, CT19 is shown as confirmation of the field classification rather than as an independent response trait. Crown transparency in 2018 (CT18) was additionally included to evaluate whether trees later classified as damaged already differed in canopy condition prior to the 2019 assessment. The number of pairs analyzed varied between 39 and 51, depending on the feature.

Crown transparency in 2019 (CT19) showed the strongest difference between healthy and damaged members of pairs and was significantly different in linear mixed models (LMMs) (***Figure 1***), with a p-value of 2 × 10^−17^. Crown transparency in 2018 (CT18), available for 39 valid pairs, was also higher in trees later classified as damaged (*P* = 2.1 × 10^−5^), indicating that these trees already showed lower canopy vitality or early drought-associated canopy deterioration in 2018. Thus, the 2019 damage classification likely reflected an amplification of pre-existing or early-emerging canopy differences rather than a completely de novo divergence.

**Figure 1.**
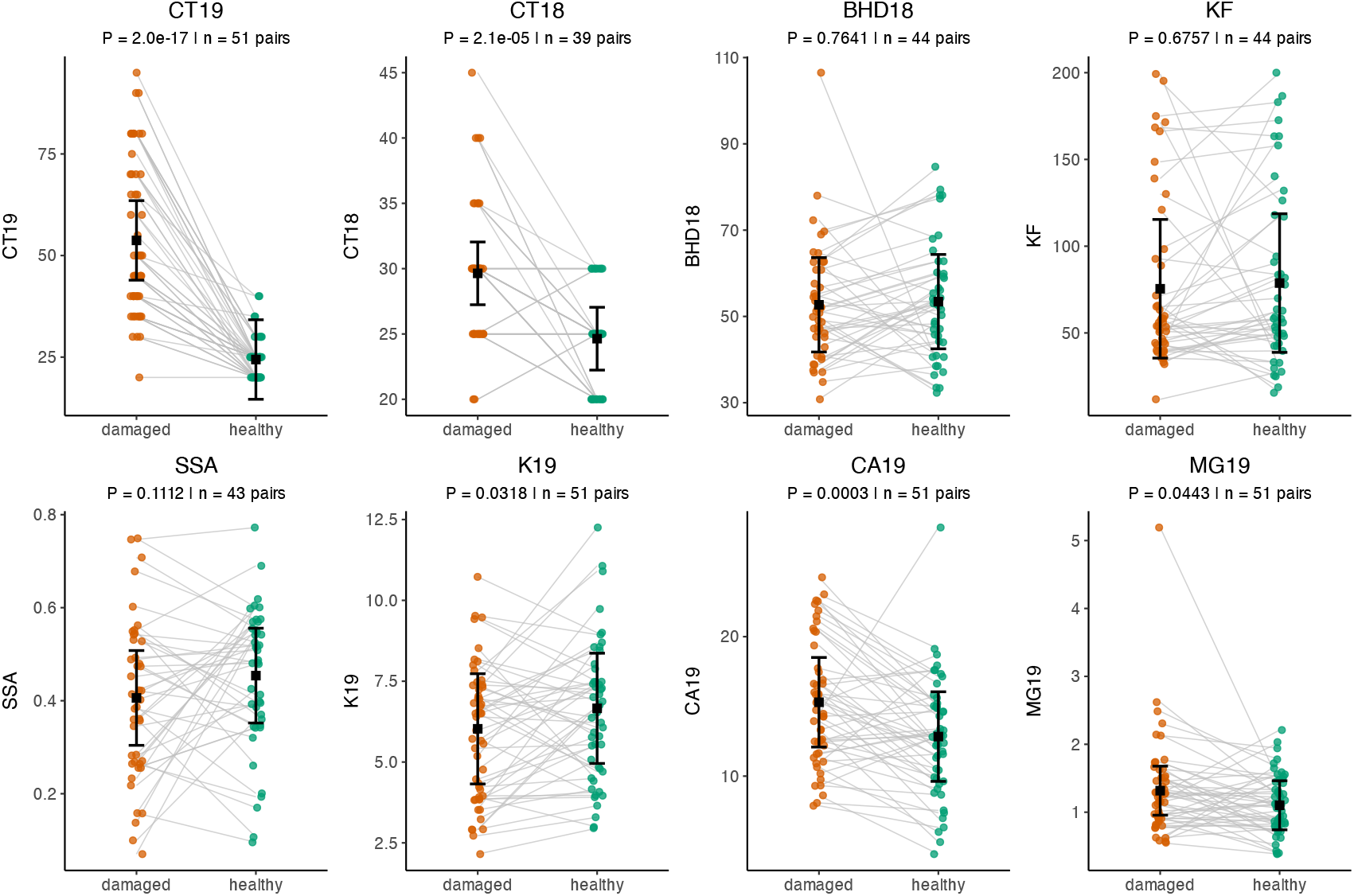
Phenotypic differences between paired healthy and damaged trees. Paired line plots display the raw data for individual tree pairs across eight assessed features: crown transparency in 2019 (CT19), crown transparency in 2018 (CT18), diameter at breast height (BHD18), crown area (KF), Safranin staining area (SSA), potassium (K19), calcium (CA19), and magnesium (MG19). Black squares and error bars represent the Estimated Marginal Means (EMMs) and 95% confidence intervals derived from the linear mixed models (LMMs). The corresponding LMM p-values and the number of pairs (*n*) are annotated above each facet.

Among the nutrient features, calcium (CA19) was approximately 20% higher in damaged members of pairs, with a p-value of 3×10^−4^. Magnesium (MG19) was slightly higher in damaged members, while potassium (K19) was slightly higher in healthy members, with p-values of 0.0443 and 0.0318, respectively. No significant differences were observed between the two groups for diameter at breast height (BHD18), crown area (KF), or other nutrients (nitrogen [N19], manganese [MN19], and phosphorus [P19], see Appendix 1–figure 1), with p-values ranging from 0.2045 to 0.7641. Interestingly, there was only a trend that safranin staining area (SSA) was higher in apparently healthy than in damaged members of a pair, although this is an indicator of xylem vessel damage due to drought. Overall, these results indicate that paired trees were broadly matched in size and crown dimensions, while trees classified as damaged showed stronger canopy damage indicators and selected physiological differences. The significant CT18 difference suggests that these trees may already have shown lower canopy vitality or early drought-associated deterioration in 2018, before the stronger divergence observed in 2019; therefore, we interpret the damage classification as drought-associated canopy decline rather than as evidence for a single exclusive causal mechanism.

### Leaf spectral signatures of drought damage

Leaf reflectance spectra of 33 healthy and 33 paired damaged trees from Baselland are shown in ***Figure 2***. Panel (a) presents spectra from the fresh leaves measured shortly after harvest, and panel (b) presents spectra from leaves dried for 72 h at 40 °C. A hierarchical spectral clustering with parallel analysis (HSC-PA) method (***Li et al., 2026***) was applied to both fresh and dried leaves, segmenting the spectra into 24 and 26 distinct patterns, respectively (Appendix 2–figure 1, Appendix 2–table 1). This technique provides a representation of spectral features while accounting for correlations among wavelengths and reducing dimensionality. In HSC-PA, “levels” refer to the depth of recursive spectral segmentation, with higher levels representing finer subdivisions of correlated wavelength regions. The final retained segments occurred at levels ranging from 2 to 5 for fresh leaves and from 3 to 6 for dried leaves, indicating differing depths of spectral similarities among the segments and distinct patterns in fresh and dry samples. Segments that retained a single principal component (PC) (Appendix 2–figure 1b, d) covered the entire spectrum from 400 to 2500 nm.

**Table 1.** Candidate genes overlapping with both top 1,000 SNPs and top 1,000 k-mers within a ± 200 bp window.

| Gene ID | Chr. | Position | Description | GO Terms |
| --- | --- | --- | --- | --- |
| Bhaga_1.g4024 | Bhaga_1 | 41,791,173–41,793,828 | benzyl alcohol O-benzoyltransferase-like | transferase activity (GO:0016747) |
| Bhaga_8.g2046 | Bhaga_8 | 17,144,517–17,148,868 | RNA polymerase II transcriptional coactivator KIWI-like | DNA binding (GO:0003677), regulation of transcription, DNA-templated (GO:0006355) |

*Note:* Three additional cross-validated SNPs (Bhaga\_1.50346833, Bhaga\_2.55973907, and Bhaga\_3.20123784) were located in intergenic regions.

**Figure 2.**
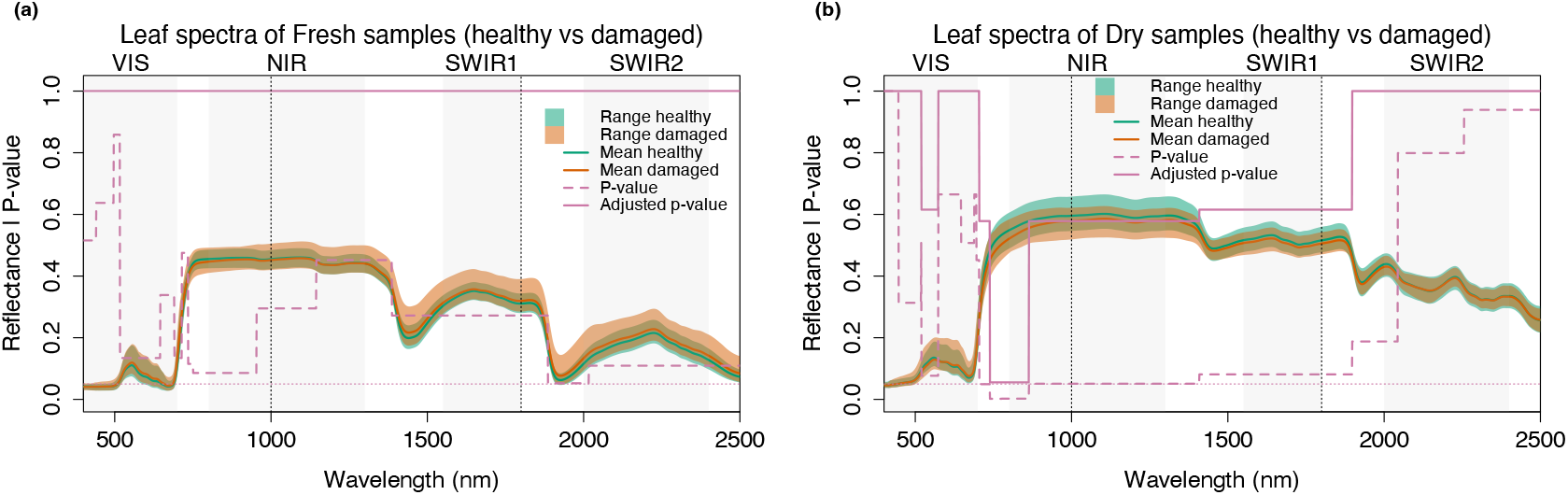
Comparison of leaf spectra across segments. (a) for fresh leaves and (b) for dried leaves. The green and orange shaded areas show the range of the leaf spectra of healthy and damaged samples, respectively. The solid green and orange lines show the mean reflectance. The pink dashed line shows the p-value from linear mixed models, while the pink solid line shows the p-values after adjustment for multiple testing.

Segmented spectral features were compared between undamaged and damaged groups using linear mixed models (***Figure 2***). In fresh samples, the minimum raw p-value (0.05) was observed in segment 15 (2017–2500 nm), though all segments yielded adjusted p-values of 1.0. Conversely, dry samples exhibited raw p-values below 0.05 across the 705–1408 nm range, encompassing segments 22, 23, 25, and 26. Following adjustment, segments 25 (739–776 nm) and 23 (777–863 nm) remained significant. LMMs conducted at individual wavelengths showed consistent patterns, albeit with reduced statistical power (Appendix 2–figure 2).

Overall, the region spanning the red light and the initial segment of the near-infrared (NIR) showed the greatest contribution to distinguishing leaves from damaged versus undamaged trees. Dry leaf spectra were consistently more informative than fresh spectra in these comparisons (***Figure 2***), indicating that the strongest differences were related to dry matter composition and not water content.

### Population structure and micro-environmental confounding

For low-coverage sequencing, we performed single-end sequencing on 275 samples in the first run, and 222 samples in the second run. The third run comprised paired-end sequencing on 173 samples. Following raw data processing, as described in Section Individuals, 244 samples remained after filtering out those with a mean coverage lower than 0.1× across the 12 chromosomes. The mean coverage of the remaining samples ranged from 0.13× to 1.16×. SNP calling, conducted with ANGSD, identified 1,261,711 SNPs. The plot of LD *r*^2^ against the distance from the focal SNP on each chromosome (Appendix 3–figure 1) showed an LD decay half-life at around 90 bp, which we applied for LD pruning. The pruned SNP set included 927,457 SNPs. PCA and Admixture analysis were performed on the LD-pruned SNP set to investigate population structure.

As shown in the PCA plot (***Figure 3***a), the first principal component (PC1) explained 2.33% of the variance, while the second principal component (PC2) accounted for 0.47% of the variance. Each tree is represented as a point on the plot, color-coded by region and shaped according to health status. The 95% confidence intervals, depicted by ellipses, indicate the genetic cohesion of trees within regions and the distinction between regions. While the variance explained by the leading PCs was low, as expected for genome-wide SNP data with many weakly correlated markers, the leading axes captured geographic structure rather than separation by health status, indicating that healthy and damaged trees originate from common genetic populations. The plot shows a separation along PC1 for samples from Birsfelden, indicating genetic differentiation between this site and other sites. Consistent with these individual-level patterns, a PCA of allele frequencies across the 18 sequenced pools also showed a separation along PC1 for pools 1 and 2, which contained the majority of Birsfelden samples (Appendix 3–figure 2).

**Figure 3.**
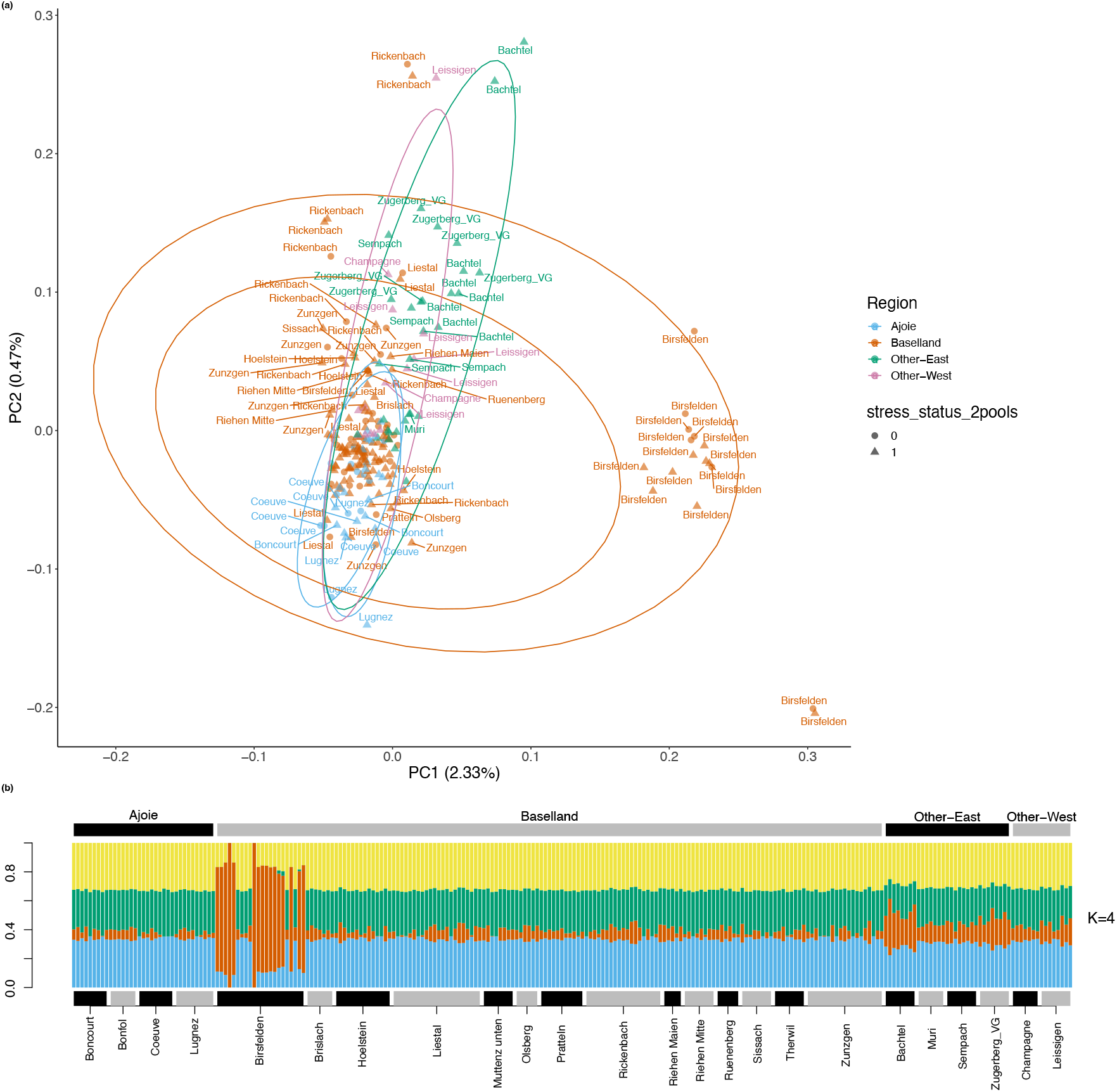
Population structure revealed by PCA and Admixture analyses on LD-pruned SNPs from low-coverage individual sequencing data. (a) PCA plot. Each point represents a single tree, labeled with site names. The x-axis and y-axis represent the first and second principal components. Regions are color-coded: Ajoie (blue), Baselland (orange), East (green), and West (pink), with health status denoted by point shape; circles for damaged trees (health status 0) and triangles for healthy trees (health status 1). Ellipses represent 95% confidence intervals for each region, with a notable separation along PC1 for most of the trees from Birsfelden. (b) Admixture analysis results for *K* = 4. Each individual tree is represented by a vertical line, partitioned into colored segments indicating estimated membership fractions in ancestral populations. The regions (top) and sites (bottom) of the samples are annotated and alternate in colour (black and grey).

***Figure 3***b presents the results of the admixture analysis for *K* = 4, and the results for *K* = 2 to *K* = 5 are shown in Appendix 3–figure 3. The corresponding log-likelihood values for each *K* are as follows: *K*_2_ = −908, 788, 006.53, *K*_3_ = −908, 747, 341.65, *K*_4_ = −908, 731, 835.25, and *K*_5_ = −908, 731, 928.13. The highest log-likelihood was obtained for *K* = 4, which also matched the geographical configuration of the four regions: Ajoie, Baselland, East Switzerland, and West Switzerland; therefore, we focus on interpreting the results for *K* = 4. The samples from Birsfelden exhibit a substantial proportion of one ancestral component (depicted in orange at *K* = 4). Additionally, the samples from the East and West regions display a larger proportion of the same ancestral component.

The observed genetic cohesion within geographic regions—and the distinct genetic profile of the Birsfelden site—co-occurred with a localized, interspersed distribution of undamaged and damaged trees. This population structure highlighted the necessity of accounting for demographic and spatial confounding in the subsequent genome-wide association studies to mitigate false-positive signals driven by local stratification.

### Genomic signatures of drought response

#### P-value inflation and genomic control

We conducted genome-wide association studies (GWAS) on the 12 pooled samples to identify genetic variants associated with damage. Because of the genetic structure in the dataset, and the available reference genome being for an out-of-dataset beech tree from central Germany that is likely more closely related to some of these samples than to others, we used both reference genome-based and reference-free approaches. For the reference-based approach, Cochran-Mantel-Haenszel (CMH) tests were applied to 11,671,486 SNPs. For the reference-free approach, we tested the frequencies of 31-bp *k*-mers across the pools. In both analyses, the initial QQ plots displayed a deviation from the expected 1:1 line (***Figure 4***b and d, red points), indicating an inflation of P-values (λ = 1.12 for SNPs, λ = 1.98 for *k*-mers). To account for population structure, together with likely unmeasured micro-environmental heterogeneity inherent to the field design, we applied genomic control to scale the test statistics. This adjustment reduced the P-value inflation, shifting distributions toward the null expectation, though deviations remained in the extreme tails, particularly in SNP associations (***Figure 4***b and d, blue points).

**Figure 4.**
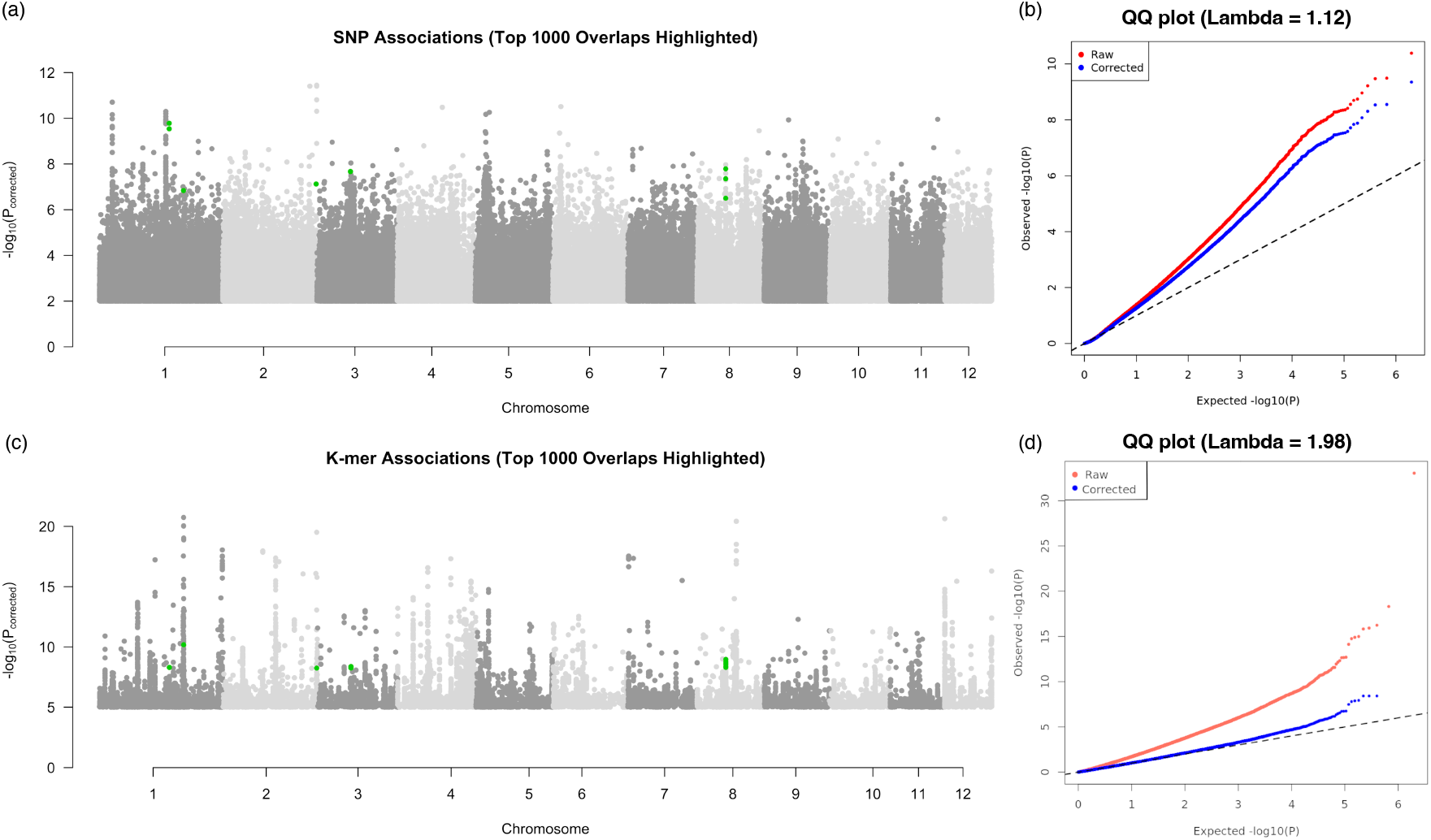
Genomic signatures of drought response and cross-method validation. (a) Manhattan plot of the reference-based SNP associations. The top 1,000 statistical outliers were compared with the top 1,000 reference-free *k*-mers, with the 8 spatially overlapping SNPs (± 200 bp) highlighted in green. (b) QQ plot for the SNP CMH test, demonstrating the severe inflation of raw P-values (red) and the effect of genomic control (λ = 1.12, blue). (c) Manhattan plot of the reference-free *k*-mer associations, highlighting the 15 overlapping *k*-mers in green. (d) QQ plot for the *k*-mer CMH test, showing extreme P-value inflation (λ = 1.98, red) corrected via genomic space transformation (blue).

#### Empirical FDR and permutation testing

Following genomic control, numerous loci retained small P-values. At an exploratory threshold of *P*_*corrected*_ < 10^−5^, we identified 5,308 SNPs and 26,462 *k*-mers. However, because traditional False Discovery Rate (FDR) adjustments assume a uniform null distribution, which was violated by our structured populations, we calculated an empirical FDR via 63 spatial permutations.

Empirical permutations indicated that background null models generated similar or higher counts of significant associations compared with the observed dataset. For both SNPs and *k*-mers, the empirical FDR consistently exceeded 1.0 across all tested significance thresholds (Appendix 4–tables 1, 2). At *P*_*corrected*_ < 10^−5^, the mean number of associations in null SNP models was 5,355 (FDR = 1.01), and for *k*-mers, the mean was 32,041 (FDR = 1.21).

#### Cross-method validation and candidate gene identification

To identify potential associations among these statistical outliers independently of absolute P-value thresholds, we extracted the top 1,000 loci from both the reference-based (SNPs) and reference-free (*k*-mers) analyses and performed a spatial cross-validation.

When intersecting the genomic coordinates of the top 1,000 *k*-mers with the top 1,000 SNPs using a ± 200 bp window to account for localized linkage disequilibrium, 8 SNPs and 15 *k*-mers were identified in both datasets (highlighted as green dots in ***Figure 4***a and c). Functional annotation of the genomic regions encompassing these eight SNPs revealed that five mapped to, or were in close proximity (± 200 bp) to, two distinct protein-coding genes (***Table 1***).

The first candidate, *Bhaga_8*.*g2046*, is annotated as an RNA polymerase II transcriptional coacti-vator (KIWI-like). The second candidate, *Bhaga_1*.*g4024*, encodes a benzyl alcohol O-benzoyltransferase-like protein. The remaining three overlapping SNPs were located in intergenic regions.

#### Absence of overlap with reported drought-associated loci

To test the universality of drought-associated genetic markers reported for beech, we further intersected our top 1,000 loci with the 76 significant drought-associated SNPs previously reported in an independent European beech cohort by Pfenninger et al. (***Pfenninger et al., 2021***). We found no spatial overlap—neither exact matches nor overlaps within a ± 200 bp window—between these previously reported SNPs and our top outliers from either the SNP or *k*-mer datasets.

## Discussion

In this study, we integrated genomic data with detailed phenotypic analyses—including canopy characteristics, nutrient contents, and leaf spectra—to investigate differences in apparent tree damage in monitored European beech stands following a severe drought event. We observed a phenomenon of patchy crown dieback in mature beech trees at Swiss forest sites, similar to observations in German cohorts (***Pfenninger et al., 2021, 2024***). While this German study identified SNPs predictive of beech performance post-drought, we hypothesized that the geographic structure of genetic variation and the high plasticity of beech trees (***Gárate-Escamilla et al., 2019***) might complicate such predictions across different regions where beech occurs. Our findings are consistent with this hypothesis: although phenotypic data clearly distinguish damaged from undamaged trees, our genomic analyses indicate that micro-environmental or historical factors may be stronger predictors of tree dieback in these populations than a universal set of genetic predictors.

### Contrasting Genomic Signatures with Other Cohorts

A primary objective of our study was to test whether genetic predictors of drought dieback could be universally applied across European beech populations. Pfenninger and colleagues(***Pfenninger et al., 2021***) previously identified 106 SNPs significantly associated with drought resistance using Pool-GWAS in a German cohort. In a recent correction, they demonstrated that using a nonparametric machine learning algorithm (eSPA*) on 20 informative loci allowed for an 88% prediction probability of the drought phenotype within their validation sample (***Pfenninger et al., 2024***).

To test the universality of these findings, we cross-referenced our top 1,000 statistical outliers from both reference-based (SNPs) and reference-free (*k*-mers) genetic association tests for post-drought damage with the published loci (***Pfenninger et al., 2021, 2024***). We found no spatial overlap between our genomic outliers and the previously reported loci. This lack of overlap suggests that European beech may not rely on a fixed, universal set of major-effect alleles to cope with drought. Instead, the genetic architecture of the drought response appears to be more context-dependent and population-specific. The predictive loci identified by Pfenninger et al. (***Pfenninger et al., 2024***) may be effective within the specific genetic background and local environmental context of their Hessian cohort, but our results indicate these specific associations do not translate to predicting different post-drought outcomes in the geographically and likely genetically distinct Swiss populations analyzed here.

In designing the study, we endeavored to work with regenerating managed forest stands known to be of local origin. However, when interpreting the genetic structure, we found that the forest in Birsfelden originated from beech trees planted in 1872-1935 by the local community (***Muttenz et al., 2009***), although this had not been previously known to monitoring organizations or to the responsible forester. The origin of the planted beech trees is yet unknown.

### Micro-environmental Variation and Phenotypic Plasticity

The lack of universal genetic predictors in our study may be further explained by the influence of local population structure. Our low-coverage sequencing revealed that even geographically proximate stands (e.g., Birsfelden) can be genetically distinct (***Figure 3***). When evaluating the pooled genomes across these structured populations, we observed substantial P-value inflation. Despite applying genomic control, our permutation tests revealed that null datasets produced similar numbers of significant associations as the true dataset (FDR > 1).

This statistical noise indicates that in our cohort, the observed condition of the trees one year after a severe drought is a complex trait with underlying contingencies that are not well accounted for in our measures of genetic variation. Rather than major-effect allelic variation, tree performance during drought may be strongly influenced by micro-environmental heterogeneity (e.g., localized soil depth or hydrology), plant–plant competition (a neighbor depriving a focal tree of water supply), and other prior or current environmental factors (***Klesse et al., 2022; Rieder et al., 2026***). The few common genetic signatures identified through *k*-mer and SNP cross-validation point towards non-genetic reasons for the differences between damaged and undamaged trees. Furthermore, the top candidate, *Bhaga_8*.*g2046*, encodes an RNA polymerase II transcriptional coactivator (KIWIlike), a regulator involved in transcriptomic reprogramming. Similarly, *Bhaga_1*.*g4024* is involved in benzenoid and phenolic glycoside biosynthesis as part of environmentally responsive or induced specialized metabolism. Because genetically similar trees may behave differently under stress due to such plastic transcriptional responses (***Gárate-Escamilla et al., 2019***), static genomic markers may have limited power to predict damage status in a heterogeneous field setting.

### Physiological and Spectral Indicators of Stress

While universal genomic predictors proved difficult to identify, our phenotypic characterization provided robust indicators of tree stress. In structural dimensions, paired trees were similar in diameter and crown area, supporting the pairing design. Contents of most measured nutrients were also similar, indicating that paired trees were similar except for damage state. However, there were differences in measured potassium and calcium content between paired apparently healthy and apparently damaged trees. These cannot be interpreted as causal because foliar nutrient concentrations were measured after drought damage had developed. Higher potassium in apparently healthy trees may have supported better stomatal regulation and turgor maintenance under water limitation, whereas lower potassium in damaged trees may indicate reduced capacity to maintain water balance (***Benlloch-González et al., 2008; Sardans and Peñuelas, 2021***). Calcium was higher in damaged trees, consistent with stress-related signaling processes, including stomatal regulation (***Knight et al., 1997; Aliniaeifard et al., 2020***). The lack of a significant difference in SSA suggests that visible canopy damage was not accompanied by a clear difference in currently functional xylem at the time of sampling. One possible interpretation is that damaged trees reduced water loss through leaf senescence or crown dieback before persistent xylem dysfunction became detectable, although a single post-drought measurement cannot reconstruct the full history of hydraulic stress.

Furthermore, the spectral data obtained from a subset of these trees provided an informative indicator of physiological decline. Using the HSC-PA method, we found that reflectance properties in the red edge and the initial NIR regions were reliable indicators of leaves sampled from visibly damaged canopies (***Figure 2***). These areas were most different in both fresh and dried leaves, but the differences had greater statistical significance in dried leaves, indicating that elimination of overlapping water absorption features rather than pigment breakdown during drying (which is, however, visible when comparing the VIS and NIR regions between panels a and b in ***Figure 2***) allowed more sensitive detection of differences that were also present in fresh leaves. The red edge is sensitive to leaf internal structure, pigment content and relative pigment degradation (***Jacquemoud and Ustin, 2019***). Spectra from dried leaves were consistently more informative than those from fresh leaves, as drying reduces the strong water absorption bands in the infrared regions, thereby revealing the underlying structural and biochemical degradation (***Carter, 1991***). This indicates that leaf discoloration in damaged canopies was distinct from any difference in leaf water content, likely resulting from a legacy of reduced water content or earlier senescence. Together with the absence of a significant SSA difference, these spectral results suggest that visible canopy decline was more clearly expressed in leaf-level structural or biochemical properties than in currently functional xylem staining measured at one post-drought time point.

### Methodological Limitations and Future Outlook

Our integrative approach highlights several methodological challenges inherent to ecological surveys of genetic associations. Sampling was conducted directly from the stressed and healthy paired trees at the time other phenotypes were sampled and recorded to ensure the direct linkage of genetic and phenotypic measures. However, this meant that DNA extracted from severely drought-stressed, top-of-canopy leaves was of inconsistent quality. We utilized a combination of low-coverage individual sequencing and Pool-seq to test scalable sequencing strategies for such cohorts. Although the low-coverage data ultimately yielded very low depth due to DNA quality issues, it was sufficient to reliably assess population structure, which proved important for interpreting results from pooled samples of extracts obtained using an optimized extraction that improved DNA quality.

Because our pooling strategy was designed to account for the regional sampling, case-control pairing and DNA quality, it mixed different population structures occurring within a region. While a paired case-control design should theoretically control for stratification, the genetic differences among populations even within one region of Switzerland likely still influenced the results, contributing to the observed P-value inflation. We were better able to mitigate this inflation using a reference-free *k*-mer approach than using reference genome-based SNP-calling, highlighting the utility of *k*-mer GWAS in avoiding reference bias in structured populations.

We note that in our study, the extracts we used for pooling followed an optimized procedure to obtain higher-quality DNA after low-coverage sequencing conducted on previous extracts of the same samples. We then used measures of DNA quality and concentration to compose pools of equal amounts of DNA from samples of similar quality and size distribution, within the bounds of the regionally structured and paired study design, similar to the pooling procedure reported by Pfenninger and colleagues (***Pfenninger et al., 2021***). However, we did not test, using low-coverage sequencing of the same samples, whether similar amounts of sample resulted in similar reads for each sample used for a pool, as has been recommended by others. Pooling is a cost-saving measure (which was also the case in our study) and if sufficient funds are available for a pre-pooling test of reads per sample from low-coverage sequencing, it may often be better to proceed with low- to mid-coverage sequencing of individual samples (***Lou et al., 2021***). Once samples are pooled, we can never be sure which reads originated from which sample in the pool, and thus we cannot be sure of even representation, although with high coverage of equimolar pools, representative allele frequencies are expected (***Rellstab et al., 2013***). Finally, pooling as a cost-saving measure is most commonly done by pooling tissue prior to extraction, which makes even representation dependent on the homogeneity of initial sample combination and extraction from the mixed-tissue sample, which can also not be verified. In our study, we dealt with the uncertainty of even representation in pooled samples together with unexpected local population structure using genomic control and permutation testing. Finally, we found the strategy of including high-coverage sequencing of selected individuals, as reported by Pfenninger and colleagues (***Pfenninger et al., 2021***), to be helpful for variant verification; as library preparation is often the larger portion of sequencing costs, this approach is less expensive per sample than a combination of low-coverage sequencing of all samples plus high coverage of (low coverage sequencing-calibrated) pools.

Overall, our results do not exclude the presence of standing natural variation for drought resistance in European beech. However, we did not find genetic variants to be reliable, standalone predictors of tree performance in northwestern Switzerland following a severe drought event using two different methods for variant detection. Future studies should sample across multiple populations with higher within-population replication to determine which aspects of genetic associations are population-specific and which are universal (***Gloss et al., 2022***). In the meantime, direct physiological indicators, including foliar nutrient status, leaf spectral traits, and canopy transparency and discoloration, remain valuable to achieve a robust approach for monitoring and predicting forest health under climate change (e.g. (***Braun et al., 2021***)).

## Conclusion

We employed a paired case-control design in monitored European beech stands to evaluate the phenotypic and genomic predictors of tree damage following a severe 2018 drought. Our phenotypic evaluations demonstrated that apparent tree drought damage is accompanied by distinct physiological and spectral signatures. Specifically, elevated leaf calcium content emerged as a consistent indicator of canopy damage, likely reflecting active stress signal transduction. Furthermore, leaf reflectance in the red edge and near-infrared regions provided an informative readout of physiological decline. Spectra from dried leaves were particularly effective in distinguishing health status by unmasking structural and biochemical degradation otherwise obscured by water absorption. In contrast, a measure of functional xylem area did not reliably differentiate undamaged from damaged trees.

Our genomic analyses indicated that damage during this extreme drought was not determined by a universal set of major-effect resistance alleles. Low-coverage individual sequencing revealed surprising population stratification within a relatively small geographic region, which likely influenced the association models and contributed to P-value inflation even though the genetically differentiated group was evenly distributed between healthy and damaged samples. Because a genomic control approach meant to account for population structure and environmental variation did not fully eliminate the observed P-value inflation, we furthermore conducted permutation testing. This permutation testing showed that random assignment of pools to healthy or damaged status generated statistical associations comparable to those in the observed dataset, meaning that the genetic associations we observed with the damaged state of trees were not robust.

We nevertheless investigated genomic outliers that were commonly identified through cross-method validation (reference-based SNPs and reference-free *k*-mers). These showed no spatial overlap with predictive loci previously reported in an independent German cohort, nor did the top 1000 strongest statistical associations initially identified using the genomic control correction with either the SNP or kmer approach (***Pfenninger et al., 2021, 2024***). This lack of cross-cohort replication, combined with the variable results of association tests, suggests that the genetic architecture of drought response in European beech is population-specific. The few outlier loci we did repeatedly identify in methodological cross-validation point toward large-scale transcriptional regulation and secondary metabolism, but these loci should be interpreted as hypothesis-generating rather than validated causal markers. Overall, our results suggest that tree performance is unlikely to be explained by a universal set of major-effect loci alone and may instead reflect population-specific, environment-dependent, and potentially polygenic drought responses.

These findings highlight the methodological challenges of conducting genome-wide associations in natural, highly structured forest populations. While standing natural variation for drought resistance likely exists within the European beech *Fagus sylvatica*, our results indicate that genomic data alone may not currently provide reliable, standalone predictions of tree performance across diverse geographic regions. Future research should prioritize high-replication sampling within discrete populations to disentangle local adaptations from universal stress responses. In the context of escalating climate change, direct physiological indicators—such as foliar calcium concentration and canopy transparency and discoloration—remain a robust approach for monitoring forest health.

## Methods and Materials

### Study Sites, Sample Collection, and Measurements

Our analysis focuses on 68 pairs of healthy and damaged European beech trees (*Fagus sylvatica*), as assessed by crown transparency, across nine monitoring sites in northwestern Switzerland: six in Baselland and three in Ajoie. The pairs were selected to represent a large range of environmental conditions such as geology and within pairs, trees were matched for age, size, position, and visible geologic conditions within the forest stand. Each pair consisted of one tree showing signs of damage and a neighbor (separated by at most one other tree) that appeared healthy based on ground assessments of crown transparency in July 2019. In addition, a subset of trees were also assessed for crown discoloration, leaf nutrient concentration, shoot growth, fructification, and proportion of active vessels, providing a holistic assessment of the trees’ health and growth condition (see Section Phenotypic analyses).

In addition to the paired samples, the study included 26 other monitored trees from the nine sites and 108 monitored trees from 14 further sites, spanning over 200 km across Switzerland. All these additional trees were apparently healthy based on crown transparency assessments. These samples were used to assess the genetic structure of the pairs in the context of Swiss beech forests (see Section Genetic analyses). ***Figure 5*** shows an overview of the study sites and their location.

**Figure 5.**
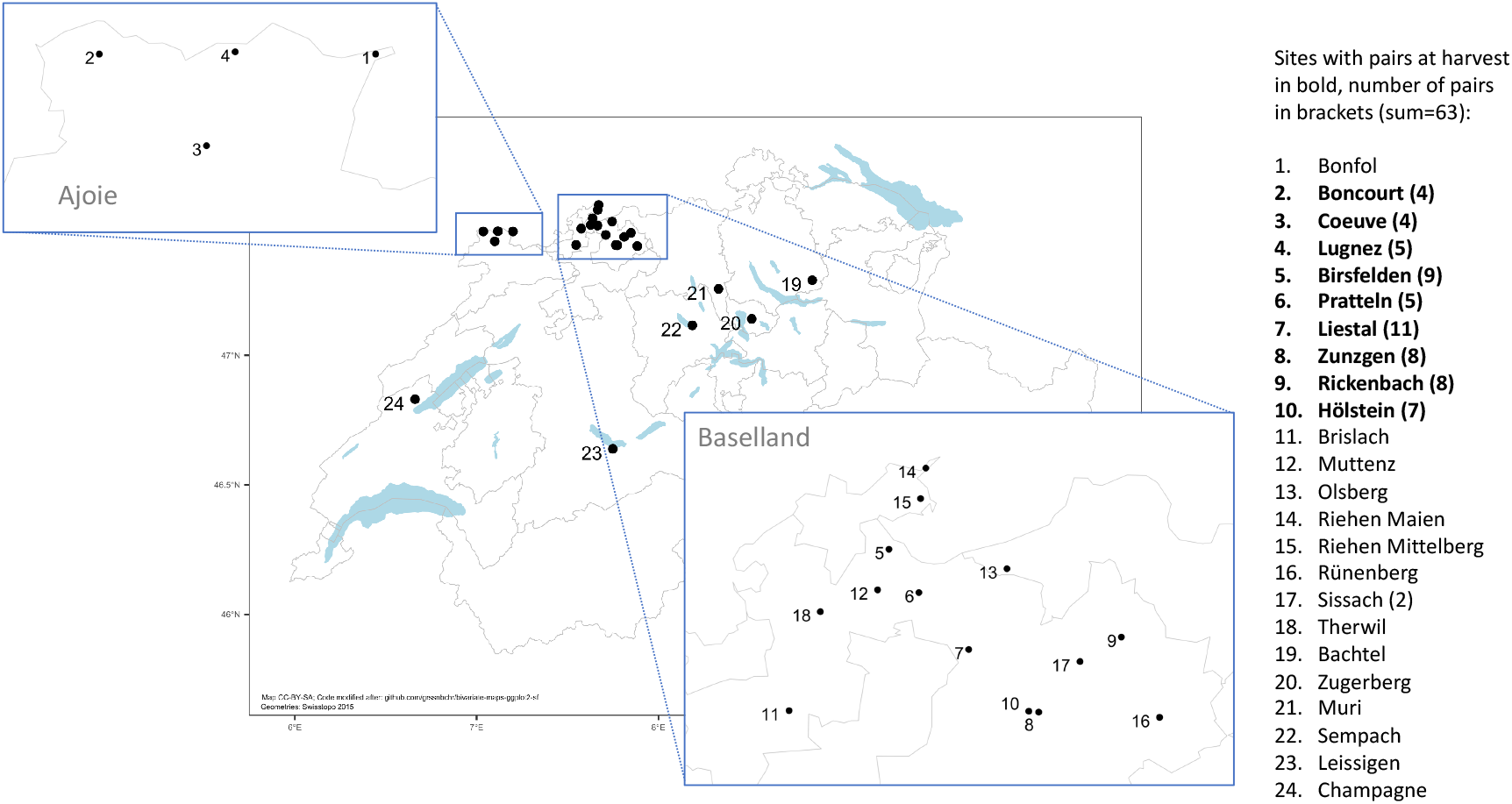
Location of study sites in Switzerland.

#### *In situ* assessment of tree and canopy status

Crown transparency and discoloration were assessed according to the recommendations of ICP Forests (***ICP Forests, 2022***) and as outlined in (***Braun et al., 2021***) between mid July and mid August. One third of the trees was assessed twice to determine repeatability.

#### Branch harvest

Most sample collection was conducted during summer 2019, when 1 m to 2 m branches from the top of the canopy of selected trees were harvested using a helicopter. This method ensured minimal damage to the surrounding environment and the trees themselves.

#### Growth and fructification

The harvested branches were also assessed for shoot growth, discoloration, and fructification. Fructification was quantified by counting the number of fruits (current year) or fruit scars (previous years) per short shoot. The diameter at breast height (DBH) at marked points was measured in 2024 and 2018. Growth was calculated from the difference of these two measurements. Crown area was measured by measuring four perpendicular crown radii.

#### Leaf spectroscopy

Following the harvest, optical measurements were promptly conducted on three leaves from each of 75 trees (a subset of the 68 pairs) spanning five sites in Baselland using a FieldSpec 4 spectro-radiometer (Analytical Spectral Devices, Inc., Malvern Panalytical, serial number 18140) equipped with a plant probe and leaf clip V2 attachments. Reflectance values per leaf were calculated from five measurements each of four different conditions (white background, leaf with white background, black background, leaf with black background) as previously described by (***Petibon et al., 2021***). These 75 trees included 33 pairs. The measurements were conducted on fresh leaves on-site within two hours of harvest, and on dry leaves after a 72-hour drying period at 40 ^°^C. This dual approach allowed for more robust assessment of leaf differences, as water absorption features suppress signatures of variation in non-pigment leaf components in the infrared part of the spectrum, whereas drying leads to breakdown of pigments and thus a different pigment profile than in fresh leaves (***Carter, 1991; Chen et al., 2012; Wang et al., 2020***).

#### Nutrient analysis

Leaves from the harvested shoots were dried at 70 ^°^C and ground. Foliar concentrations of nitrogen (N), phosphorus (P), potassium (K), magnesium (Mg) and manganese (Mn) were analysed as described in Braun et al. (***Braun et al., 2020a***).

#### Xylem staining

Active xylem vessels were stained by infiltrating shoots with a safranine solution under moderate pressure and the proportion of stained vessels was analysed quantitatively with an image analyser as previously described by Braun et al. (***Braun et al., 2021***).

### DNA extraction and sequencing

Leaves were sampled from the harvested branches of 270 trees (2 x 68 + 26 + 108, see Section Study Sites, Sample Collection, and Measurements) and stored on silica gel at ambient temperature until DNA extraction for genetic analysis. For the focal pairs, DNA was extracted from leaves originating from the same top-of-canopy branches used for the phenotypic analyses, ensuring the relationship of our genetic and phenotypic measures in light of possible genetic differences between branches due to somatic mutations (***Schmid-Siegert et al., 2017***). However, the samples thus represented older and sometimes stressed leaves, and extracted DNA showed some degradation and some samples had low 260/230 nm ratios.

#### Low-coverage sequencing for population structure

DNA extraction was initially conducted using the Qiagen DNeasy Plant Mini Kit (Qiagen, Hombrechtikon), modified to improve yield. Specifically, 10 mg to 15 mg of dried leaf material was utilized for each extraction. The procedure was carried out in accordance with the manufacturer’s protocol with the following modification: an additional digestion step using proteinase K was introduced following the second step of the protocol (AP1 and RNase A). During this step, 10 µL of proteinase K (10 mg mL^−1^) was added to each tube, followed by vortexing and a 15-minute incubation at 65 ^°^C. This step is supposed to increase yield by degrading nucleases and contaminating proteins, and improving lysis.

The sequencing of the extracted DNA was done at the Functional Genomics Center Zurich, using the Illumina NovaSeq platform and the TruSeq DNA PCR-Free HT Kit to reduce artifacts and bias introduced by PCR-based library preparation; however, this resulted in very low coverage from these samples, which may be due to residual inhibitors from the plant tissue. Quality control of remaining DNA samples by microvolume spectrophotometer (DS-11+, DeNovix) revealed that almost all had 260/280 nm ratios at or above 1.8, but 260/230 nm ratios below 1.8. In order to obtain as much sequence information as possible from this approach, we conducted two runs of single-end sequencing, producing reads of 101 base pairs (bp) each, and a subsequent third run of paired-end sequencing, generating reads of 151 bp.

#### Higher coverage sequencing and poolseq

We aimed to obtain higher-coverage sequences from the same samples. To this end, we switched to the Norgen Plant and Fungi genomic DNA extraction kit (Norgen Biotek, Thorold, ON, Canada) with some minor modifications, because we had optimized this procedure to obtain higher-quality DNA from dried top-of-canopy leaf samples (***Czyż et al., 2023***). Specifically, 30 mg of dried leaf laminar tissue were transferred to 2 mL tubes (Eppendorf, Hamburg, Germany) containing two 3 mm stainless metal beads and frozen in liquid N2. Then, samples were ground with a TissueLyser II (Qiagen, Hombrechtikon, Germany) adjusted at 25 Hz for 2 min. The powder was briefly centrifuged before adding 750 µL of lysis buffer and vortexed to mix with the buffer. 1 µL of RNAse A (DNAsefree, 100,000 units/ml in 50% glycerol, 10 mM Tris-HCl, pH 8.0) and 3 µL Proteinase K (>600 mAU/ml, ~20 mg mL^−1^ in 10 mM Tris-HCl pH 7.5, containing calcium acetate and 50% v/v glycerol) were added to the lysis buffer prior to 10 min incubation at 56 ^°^C with 1000 agitation. Then an additional 1 µL of RNAse A was subsequently added to the reaction, followed by a 10 min incubation at 65 ^°^C with 2000 agitation. Next, 150 µL of Binding Buffer I was added to each tube, thoroughly vortexed, and incubated for 5 min on ice. The remaining steps of the extraction procedure were carried out following the manufacturer’s protocol with the modification that 70% ethanol was mixed with lysate by pipetting ca. 10x/sample instead of vortexing. Finally, the purified DNA was eluted in 100 µL of elution buffer and stored at −20 ^°^C. The quantification of purified DNA was performed using a fluorescence assay specific for dsDNA (Thermo Fisher Qubit, dsDNA HS Assay Kits) and analysis of purity was done by microvolume spectrophotometer (DeNovix DS-11+), while agarose gel electrophoresis was used to assess DNA degradation for pools and several individual samples.

DNA extracts were pooled based on their health status, DNA quality, and sites of origin (Supplementary file 1). We thus ensured that samples in each pool were of similar quality as assessed by spectrophotometry, and then pooled based on total amounts of dsDNA as measured by fluorescence assay (Qubit), assuming similar size distributions and given that DNA would be fragmented in the course of library preparation. Pooling was furthermore done in a paired manner, ensuring that each “damaged” tree in pool A had its corresponding “healthy” counterpart in pool B and so on. This resulted in six replicate pools (comprising six damaged and six healthy pools) and an additional six pooled regional controls from samples originating at other monitored beech forests across Switzerland that were not part of the case-control design. This allowed us to maintain the paired design while ensuring even coverage as much as possible by combining samples of similar DNA quality and as much as possible keeping regions separate in case of geographic differences in population structure. These pooled samples were then dispatched to Novogene for library construction using PCR-based library preparation and sequencing. Sequencing was conducted using the Illumina NovaSeq 6000 platform, employing a 150 bp paired-end sequencing approach.

In addition to the pooled samples, 14 individual trees were chosen for high-coverage sequencing based on DNA quality. This optimized the use of available positions on the sequencing lanes and allowed a more in-depth analysis of the candidate genes identified through Pool-GWAS.

### Phenotypic analyses

Data processing and analyses were conducted using R (***R Core Team, 2023***), with the majority of plots generated using the ggplot2 package (***Wickham, 2016***). The scripts for these analyses are also provided (see Data and Code Availability).

#### Monitoring dataset

The monitoring dataset was provided by the Institut fuer angewandte Pflanzenbiologie (IAP) according to published protocols (***Braun et al., 2021***) and analyzed to support the evaluation and interpretation of differences between the paired healthy and damaged trees. In the initial phase of data preparation, features with more than 40% missing data were excluded, except for crown transparency in 2018 (CT18), which was retained a priori as a pre-2019 canopy-condition variable. To ensure valid comparisons, only tree pairs where both the healthy and damaged trees had non-missing values for the feature of interest were included in the analysis. For each feature, valid pairs were identified, and the data were subset to retain only these pairs.

To account for the hierarchical structure of the data, including both site-level and pair-level variability, we fitted linear mixed models (LMMs) with tree health status (healthy vs. damaged) as the fixed effect and random intercepts for sites and tree pairs. This approach corrects for the nonindependence of trees within the same pair and accounts for site-specific variability, providing a reliable estimate of the effect of tree health status on each phenotype. Estimated Marginal Means (EMMs) and 95% confidence intervals were computed to provide adjusted group means. To visualize the data distribution alongside these adjusted overall differences, paired line plots of the raw data for individual tree pairs were overlaid with the LMM-derived EMMs and confidence intervals.

#### Spectral data

We analyzed the spectral range (R package spectrolab (***Meireles et al., 2017***)) from 400-2500 nm because the regions from 350-399 nm have a poor signal:noise ratio (***Cavender-Bares et al., 2016***) and relatively high measurement uncertainty (***Petibon et al., 2021***) and are commonly removed prior to downstream analysis. The first scan was excluded because of the potential influence from previous readings or the opening of the leaf clip. Then, the averaged reflectance under each condition was obtained with the mean reflectance of the last four scans. The calculated reflectance of a sample was obtained from the mean of scans above (*R*_*c*_) using the following formula from Miller et al. (***Miller et al., 1992***), based on Lillesaeter (***Lillesaeter, 1982***):

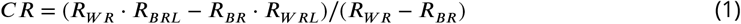

The mean leaf spectra of three leaves per tree was used for the leaf reflectance spectra of each sample.

We used the Hierarchical Spectral Clustering with Parallel Analysis (HSC-PA) method (***Li et al., 2026***) on both the fresh leaf spectra and dry leaf spectra. This generated a set of segmented features based on the correlations of wavelengths within each dataset, with each segment represented by a single principal component (PC). This approach reduces the dimensionality of the spectral data to improve power for statistical testing and accounts for autocorrelation while resulting in features that can be interpreted in terms of known trait-spectrum relationships. Subsequently, linear mixed models were applied to each spectral segment to compare healthy and damaged groups of samples. The models included random effects to account for variability across sites and paired trees, with health status as the fixed effect. To account for the multiple tests conducted across wavelengths, we adjusted the p-values employing the “BY” method as proposed by Benjamini and Yekutieli (***Benjamini and Yekutieli, 2001***). For comparison, we also performed linear mixed models on each individual wavelength.

### Genetic analyses

#### Individuals

For the low-coverage sequencing dataset, single-end reads and only the forward reads (R1) from the paired-end runs were utilized. Processing all data as single-end ensured consistency across samples and mitigated systematic biases introduced by mixing sequencing methods (***Sanchez Herrero et al., 2021***). Trimmomatic (***Bolger et al., 2014***) was used to trim adapter sequences and low-quality bases (ILLUMINACLIP:TruSeq3-SE:2:30:10, LEADING:3, TRAILING:3, SLIDINGWINDOW:4:15, MINLEN:50). Reads were aligned to the chromosome-level *Fagus sylvatica* reference genome (***Mishra et al., 2022***) using Bowtie2 (***Langmead and Salzberg, 2012***). Duplicate reads were marked and removed using Picard (***Broad Institute, (Accessed: 2018/02/21; version 2*.*17*.*8***). Samtools (***Danecek et al., 2021***) was employed to sort, merge, and filter alignments (minimum mapping quality = 20). Samples with a mean coverage below 0.1× across the 12 chromosomes were excluded.

Single nucleotide polymorphisms (SNPs) were called using ANGSD (***Korneliussen et al., 2014***) based on genotype probabilities, requiring a minimum individual count of 49, a minor allele frequency (MAF) ≥ 0.02, and a SNP p-value threshold ≤ 0.01. Linkage disequilibrium (LD) was estimated for SNPs within 1000 bp using ngsLD (***Fox et al., 2019***). LD pruning was executed with a minimum weight threshold of 0.2 and a maximum correlation coefficient of 0.9. PCAngsd (***Meisner and Albrechtsen, 2018***) was used to perform PCA and Admixture analyses on the LD-pruned SNPs, evaluating models for *K* = 2 to 5.

For individual high-coverage sequencing samples, raw data were processed using the same pipeline, except Trimmomatic was executed in paired-end (PE) mode (MINLEN:50). SNPs were called using GATK HaplotypeCaller (***Poplin et al., 2017***) with hard filtering parameters: QD < 2.0, FS > 80.0, MQ < 50.0, MQRankSum < −12.5, ReadPosRankSum < −8.0, SOR > 4.0, and QUAL < 10.0.

#### Pool-sequencing and reference-based association testing

Raw reads from the 12 pooled samples were processed using the high-coverage individual pipeline. Popoolation2 (***Kofler et al., 2011***) was utilized for SNP calling and indel filtering (5 bp window). PCA was performed on allele frequencies across all pools to assess population stratification.

To identify genetic variants associated with tree damage status, Cochran-Mantel-Haenszel (CMH) tests (***Mantel and Haenszel, 1959***) were executed across the paired pools using Popoolation2. SNPs were filtered for a minimum count of four, and coverage between 5× and 100×. To account for population stratification and environmental confounding contributing to P-value inflation, genomic control (λ) was applied (***Devlin and Roeder, 1999***). The λ inflation factor was calculated as the median of the observed χ^2^ distribution divided by the theoretical median of a χ^2^ distribution with one degree of freedom (0.4549). Raw P-values were transformed to χ^2^ statistics, scaled by λ, and converted back to adjusted P-values in logarithmic space to prevent float underflow.

Because traditional false discovery rate (FDR) methods assume a uniform null distribution of P-values, we calculated an empirical FDR via permutation testing (***Churchill and Doerge, 1994; Storey and Tibshirani, 2003***). We generated 63 null datasets by randomly permuting the “Healthy” and “Damaged” phenotype labels strictly within each paired sampling site. The empirical FDR for a given significance threshold was calculated as the mean number of significant hits across the 63 null permutations divided by the observed number of significant hits in the true dataset.

#### Reference-free *k*-mer association testing

To bypass potential reference bias and capture complex structural variations, a reference-free *k*-mer genome-wide association study was conducted. Adapting existing presence/absence *k*-mer frameworks (***Voichek and Weigel, 2020***) for Pool-seq, we used quantitative *k*-mer counts as proxies for allele frequencies.

Raw paired-end reads were decomposed into 31-mers using KMC3 (***Kokot et al., 2017***). To reduce sequencing noise, only *k*-mers appearing at least three times within a given pool were retained. The output databases were merged using `kmc_tools` to construct a unified count matrix. This matrix was subjected to three filters to remove uninformative or repetitive sequences: (1) presence in ≥ 6 out of 12 pools; (2) a minimum count of ≥ 4 in at least one pool; and (3) a maximum count threshold of 100 in any pool.

The quantitative counts of each filtered 31-mer were tested against damage status using the CMH test across paired pools. Genomic control was applied using an independently calculated *k*-mer λ factor, and empirical FDR was evaluated using the 63 within-site permutations. Statistical outliers (*P*_*corrected*_ < 10^−5^) were mapped to the *Fagus sylvatica* reference genome using Bowtie2 (`-k 1 --very-sensitive`), retaining only perfect alignments.

#### Cross-method validation and candidate gene identification

To identify robust loci independently of inflated P-values, the top 1,000 statistical outliers from both the SNP and mapped *k*-mer datasets were extracted. Using the GenomicRanges package in R, the physical coordinates of the top *k*-mers were intersected with the top SNPs using a ± 200 bp window to account for localized linkage disequilibrium.

Additionally, a spatial intersection was performed between our top 1,000 loci and 76 drought-associated SNPs previously identified in an independent European beech cohort by Pfenninger and colleagues (***Pfenninger et al., 2021***). Historical coordinates were mapped to the updated reference genome, and intersections were evaluated using the same ± 200 bp window. Finally, structural genes located within or adjacent to the loci validated by both our SNP and *k*-mer analyses were extracted from the reference GFF3 file for functional annotation.

## Data and Code Availability

All genomic and phenotypic data, as well as the analytical scripts used in this study, are available to ensure reproducibility.

- **Sequencing Data:** Raw sequencing reads (low- and high-coverage individual sequencing and pool sequencing) are available in the European Nucleotide Archive (ENA) under the study accession number **PRJEB97125** (Secondary Accession: ERP179716).
- **Sample Metadata and Pooling Information:** Tree-level sequencing metadata and ENA accessions, phenotypic measurements, DNA quality measurements, and the composition of the 18 sequenced pools are provided in Supplementary file 1.
- **Code and Derived Data:** All scripts for bioinformatic processing, statistical analyses, and figure generation are available at the following GitHub repository: https://github.com/licheng1221/Drought-susceptibility-in-European-beech-forests.
- **Permanent Archiving:** A permanent snapshot of the code will be archived on Zenodo.

## Authors’ Contributions

CRediT taxonomy roles are listed with authors in alphabetical order. Conceptualization: B.S., L.D.W., M.C.S., M.E.S., S.B. Data curation: C.L., L.D.W., E.A.C. Formal analysis: C.L., E.A.C., L.D.W. Funding acquisition: L.D.W., M.E.S., S.B. Investigation: C.L., E.A.C., L.D.W., S.H. Methodology: A.M., B.S., C.L., E.A.C., L.D.W., M.C.S. Project administration: M.C.S., S.B. Resources: S.B., M.E.S. Supervision: M.C.S., S.B. Visualization: C.L., L.D.W. Writing—original draft: C.L. Writing—review and editing: A.M., B.S., C.L., E.A.C., L.D.W., M.C.S., M.E.S., S.B., S.H.

## Acknowledgements

The authors acknowledge funding from the NOMIS Foundation; the University of Zurich, including the University Research Priority Program on Global Change and Biodiversity; the Institute for Applied Plant Biology; and U. Meier and the Baselland Forestry Office (Amt für Wald Basel Landschaft). We thank our division in the Department of Geography at the University of Zurich, the Remote Sensing Laboratories, for support and for the shared use of equipment. We are grateful to D.M. Leigh for helpful discussions regarding genetic analyses and to D. Coq–Etchegaray and L. Rohrbach for finding documentation of the historical planting of beech at the Birsfelden site. We also thank M. Ehmig for his careful reading of the manuscript and comments on the interpretation of the results.

## Appendix 1

**Appendix 1—figure 1.**
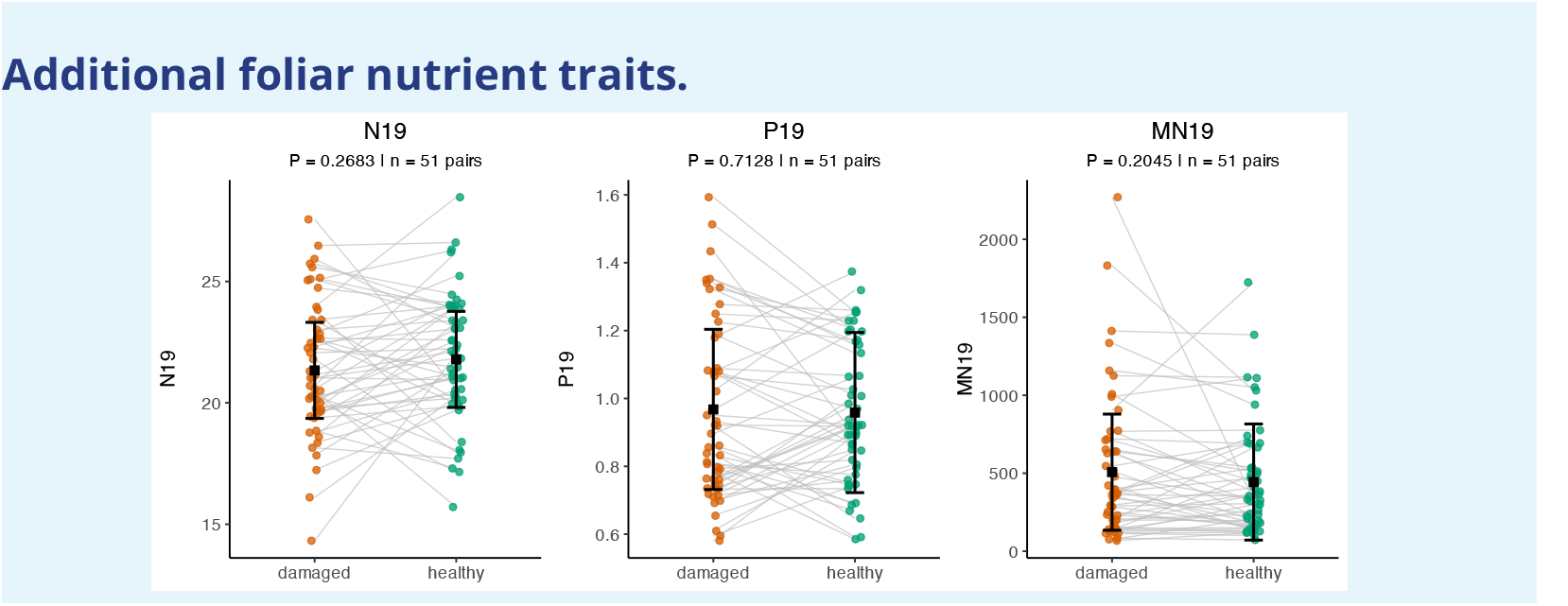
Additional foliar nutrient traits in paired damaged and healthy trees. Paired line plots show nitrogen (N19), phosphorus (P19), and manganese (MN19). Grey lines connect paired trees. Black squares and error bars represent estimated marginal means and 95% confidence intervals from linear mixed models.

## Appendix 2

**Appendix 2—figure 1.**
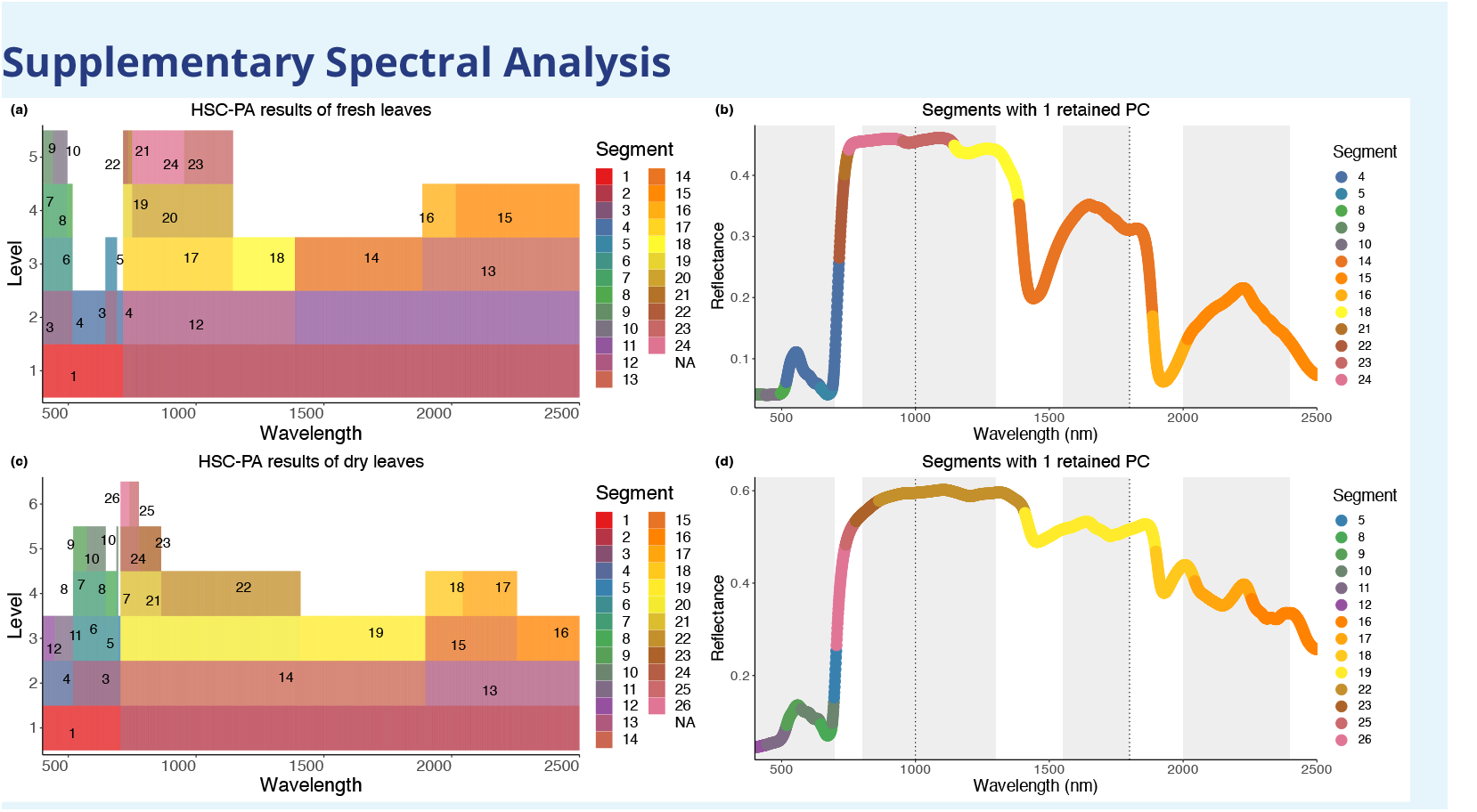
Results of Hierarchical Spectral Clustering with Parallel Analysis (HSC-PA). (a) A total of 24 segments, distinguished by various colors, were identified through spectral clustering on fresh leaves. Blank regions (NA) indicate segments that retained only one Principal Component (PC) at the previous level, preventing further clustering. (b) This panel displays the 13 segments from the fresh leaves, represented as example spectra, each associated with one retained Principal Component (PC). Panels (c) and (d) depict the results for dry leaves, which resulted in 26 distinct segments, within which 14 segments had one retained PC. The interpretation and representation in these panels are consistent with those in (a) and (b).

**Appendix 2—table 1.** Spectral range of segments identified by HSC-PA.

| Fresh Leaves |  | Dry Leaves |  |
| --- | --- | --- | --- |
| Segment | Spectral Range (nm) | Segment | Spectral Range (nm) |
| 1 | 400–714 | 1 | 400–704 |
| 2 | 715–2500 | 2 | 705–2500 |
| 3 | 400–516, 646–690 | 3 | 519–704 |
| 4* | 517–645, 691–714 | 4 | 400–518 |
| 5* | 646–690 | 5* | 696–704 |
| 6 | 400–516 | 6 | 519–695 |
| 7 | 400–496 | 7 | 520–646, 689–695 |
| 8* | 497–516 | 8* | 519, 647–688 |
| 9* | 400–440 | 9* | 520–573 |
| 10* | 441–496 | 10* | 574–646, 689–695 |
| 11 | 1387–2500 | 11* | 447–518 |
| 12 | 715–1386 | 12* | 400–446 |
| 13 | 1886–2500 | 13 | 1898–2500 |
| 14* | 1387–1885 | 14 | 705–1897 |
| 15* | 2017–2500 | 15 | 1898–2255 |
| 16* | 1886–2016 | 16* | 2256–2500 |
| 17 | 715–1144 | 17* | 2044–2255 |
| 18* | 1145–1386 | 18* | 1898–2043 |
| 19 | 715–750 | 19* | 1409–1897 |
| 20 | 751–1144 | 20 | 705–1408 |
| 21* | 735–750 | 21 | 705–863 |
| 22* | 715–734 | 22* | 864–1408 |
| 23* | 954–1144 | 23* | 777–863 |
| 24* | 751–953 | 24 | 705–776 |
|  |  | 25* | 739–776 |
|  |  | 26* | 705–738 |

Note: \* indicates segments that retained only one Principal Component (PC).

**Appendix 2—figure 2.**
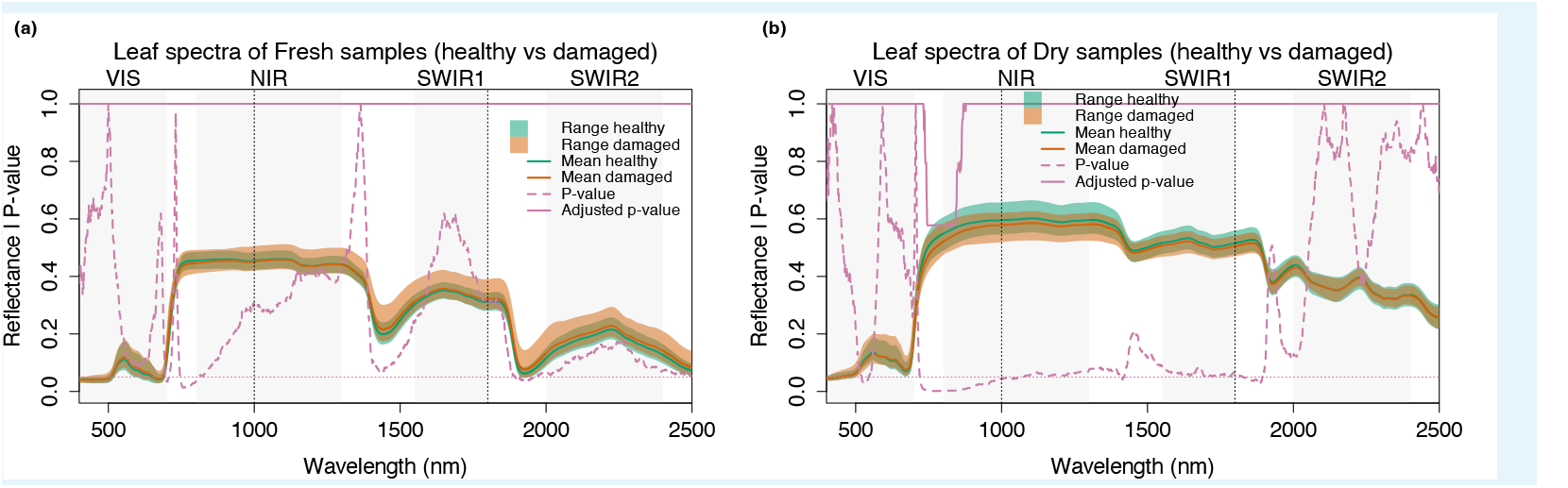
Comparison of leaf spectra at each individual wavelength. (a) for fresh leaves and (b) for dry leaves. Green and orange shaded areas show the reflectance range of healthy and damaged samples, respectively. Solid green and orange lines represent mean reflectance. The pink dashed line indicates raw p-values from linear mixed models, and the pink solid line represents p-values after Benjamini-Yekutieli adjustment.

## Appendix 3

**Appendix 3—figure 1.**
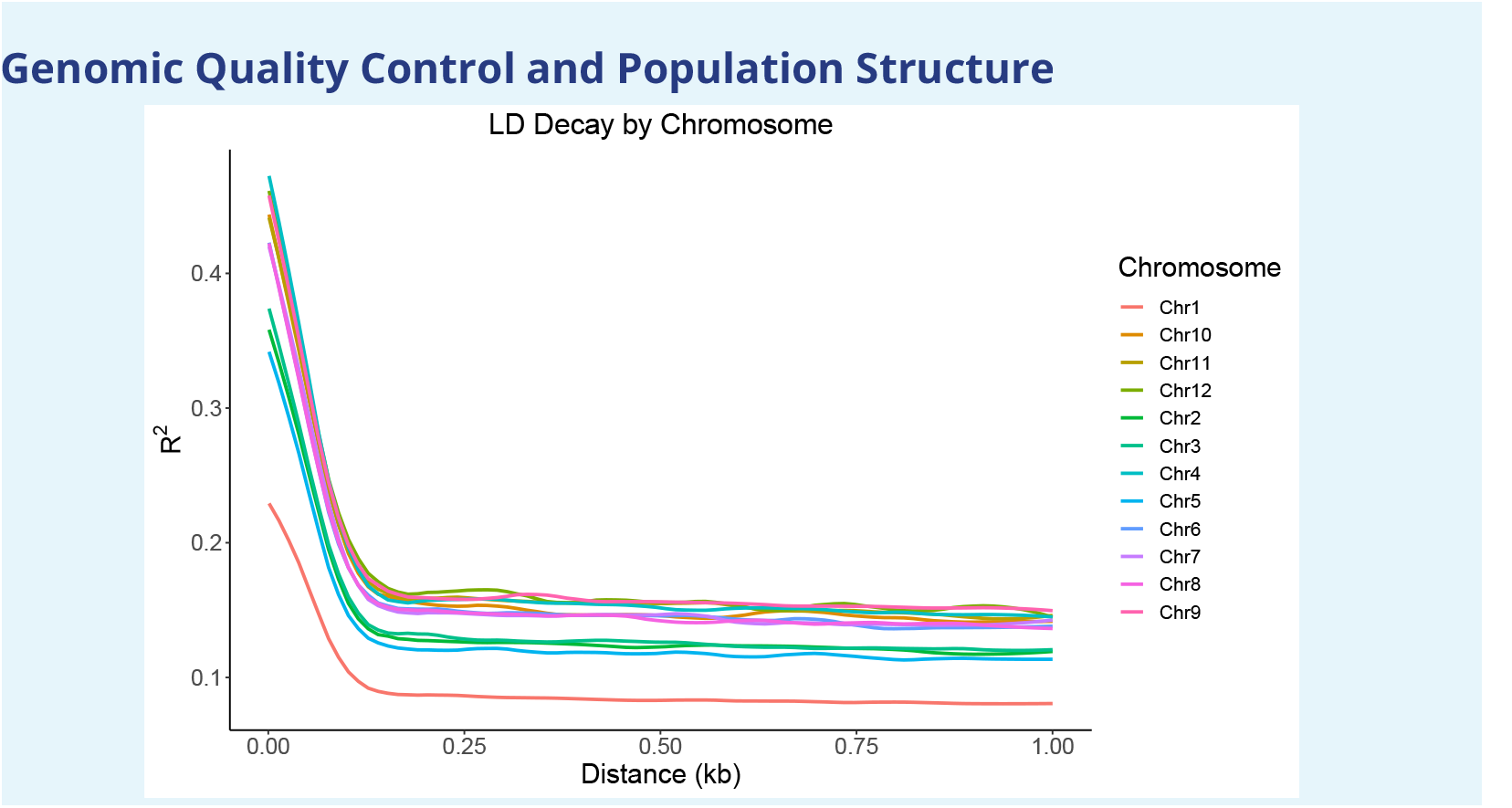
Linkage disequilibrium (LD) decay by chromosome. The decay half-life was observed at approximately 90 bp, which informed the threshold for subsequent LD pruning.

**Appendix 3—figure 2.**
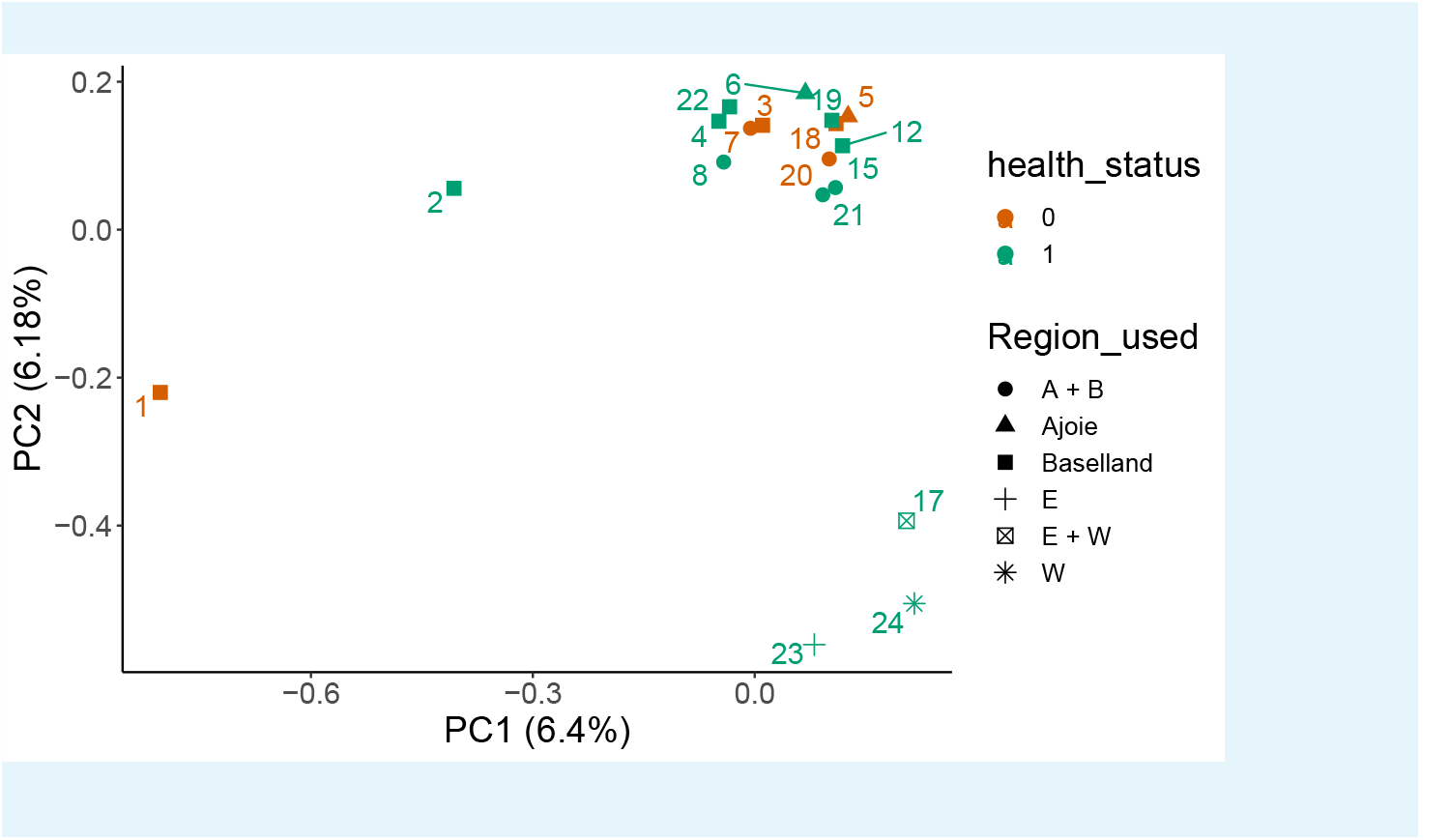
Principal Component Analysis (PCA) of allele frequencies across all 18 pools. Each point represents a pool, color-coded by health status (damaged: orange; healthy: green). Symbols indicate the geographic regions of origin. Pools 1 and 2, which contain the majority of Birsfelden samples, show separation along PC1.

**Appendix 3—figure 3.**
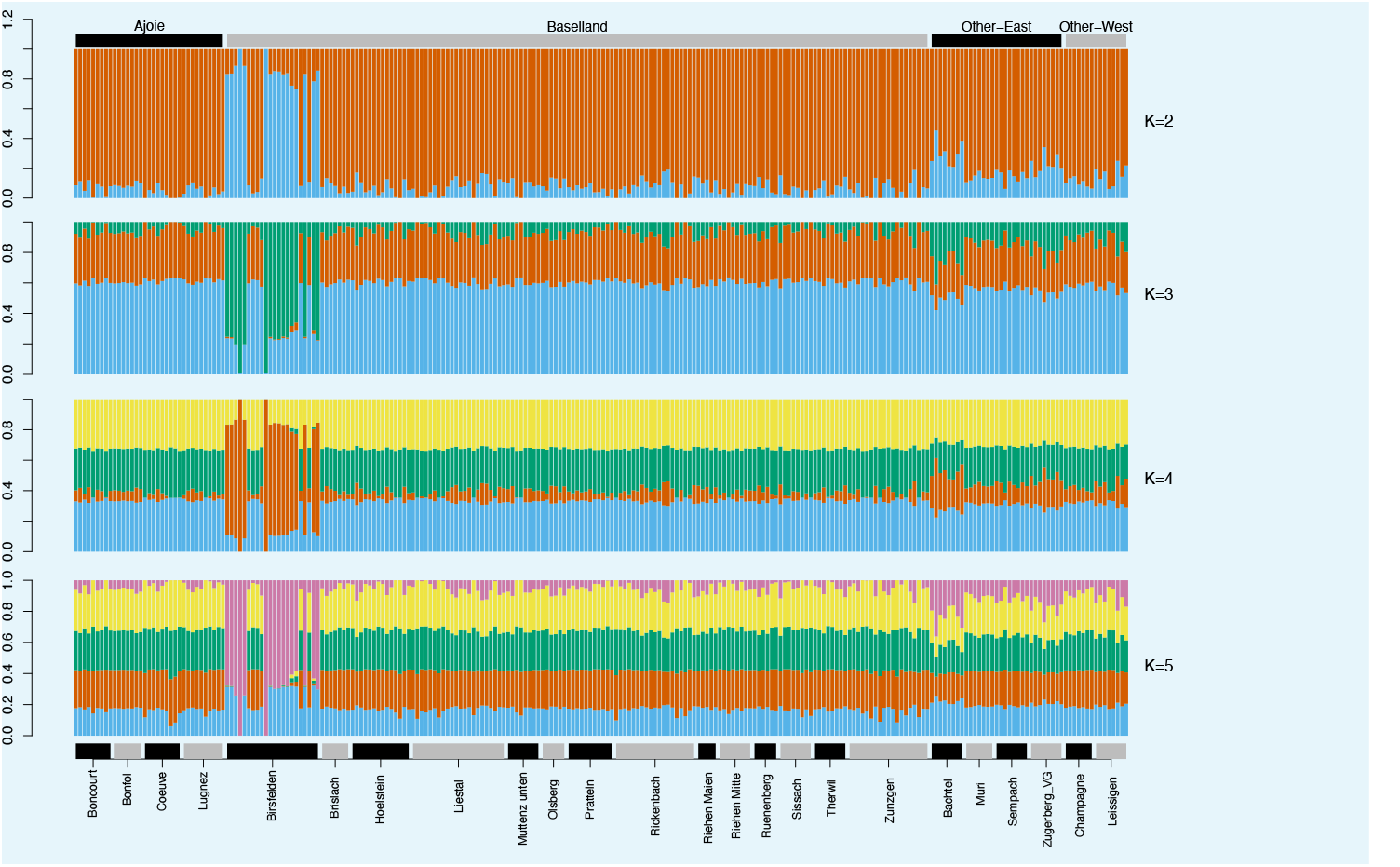
Admixture analysis results for *K* = 2 to *K* = 5 based on LD-pruned SNPs from low-coverage individual sequencing data. Each individual tree is represented by a vertical line partitioned into colored segments indicating estimated membership fractions in ancestral components. The regions and sites of the samples are annotated above and below the plot, respectively.

## Appendix 4

### Empirical FDR Estimations

The following tables present the empirical False Discovery Rate (FDR) calculated via 63 spatial permutations, where phenotype labels were shuffled within sampling sites.

**Appendix 4—table 1.** Empirical FDR for SNP-based CMH tests.

| P-value Threshold | Observed Hits | Mean Null Hits | Empirical FDR |
| --- | --- | --- | --- |
| $1 \times 10^{-4}$ | 20,231 | 20,542.7 | 1.015 |
| $3.16 \times 10^{-5}$ | 10,309 | 10,428.7 | 1.012 |
| $1 \times 10^{-5}$ | 5,308 | 5,355.6 | 1.009 |
| $3.16 \times 10^{-6}$ | 2,781 | 2,803.7 | 1.008 |
| $1 \times 10^{-6}$ | 1,437 | 1,487.7 | 1.035 |
| $3.16 \times 10^{-7}$ | 789 | 804.9 | 1.020 |
| $1 \times 10^{-7}$ | 438 | 447.2 | 1.021 |
| $3.16 \times 10^{-8}$ | 217 | 257.6 | 1.187 |
| $1 \times 10^{-8}$ | 96 | 151.0 | 1.573 |
| $3.16 \times 10^{-9}$ | 61 | 96.0 | 1.573 |
| $1 \times 10^{-9}$ | 37 | 65.2 | 1.762 |

**Appendix 4—table 2.** Empirical FDR for k-mer-based CMH tests.

| P-value Threshold | Observed Hits | Mean Null Hits | Empirical FDR |
| --- | --- | --- | --- |
| $1 \times 10^{-4}$ | 121,024 | 134,582.5 | 1.112 |
| $1 \times 10^{-5}$ | 26,462 | 32,041.5 | 1.211 |
| $1 \times 10^{-6}$ | 8,881 | 12,641.5 | 1.423 |
| $1 \times 10^{-7}$ | 4,293 | 6,855.0 | 1.597 |
| $1 \times 10^{-8}$ | 2,508 | 4,180.6 | 1.667 |
| $1 \times 10^{-9}$ | 1,634 | 2,696.7 | 1.650 |
| $1 \times 10^{-10}$ | 1,039 | 1,803.6 | 1.736 |
| $1 \times 10^{-11}$ | 704 | 1,248.4 | 1.773 |
| $1 \times 10^{-12}$ | 428 | 891.1 | 2.082 |
| $1 \times 10^{-13}$ | 268 | 638.9 | 2.384 |
| $1 \times 10^{-15}$ | 145 | 327.4 | 2.258 |
| $1 \times 10^{-17}$ | 85 | 186.6 | 2.195 |
| $1 \times 10^{-20}$ | 21 | 94.6 | 4.503 |
| $1 \times 10^{-25}$ | 1 | 28.4 | 28.429 |
| $1 \times 10^{-26}$ | 0 | 22.4 | — |

## Supplementary files

**Supplementary file 1**. Tree metadata, phenotypic measurements, DNA quality, and pooling information.

## References

Aliniaeifard S, Shomali A, Seifikalhor M, Lastochkina O. Calcium signaling in plants under drought. Salt and drought stress tolerance in plants: signaling networks and adaptive mechanisms. 2020; p. 259–298.

Asner GP, Martin RE. Spectranomics: Emerging science and conservation opportunities at the interface of biodiversity and remote sensing. Global Ecology and Conservation. 2016; 8:212–219.

Astle W, Balding DJ. Population Structure and Cryptic Relatedness in Genetic Association Studies. Statistical Science. 2009; 24(4):451–471. https://doi.org/10.1214/09-STS307, doi: 10.1214/09-STS307.

Atwell S, Huang YS, Vilhjálmsson BJ, Willems G, Horton M, Li Y, Meng D, Platt A, Tarone AM, Hu TT, et al. Genome-wide association study of 107 phenotypes in Arabidopsis thaliana inbred lines. Nature. 2010; 465(7298):627–631.

Benjamini Y, Hochberg Y. Controlling the false discovery rate: a practical and powerful approach to multiple testing. Journal of the Royal statistical society: series B (Methodological). 1995; 57(1):289–300.

Benjamini Y, Yekutieli D. The control of the false discovery rate in multiple testing under dependency. Annals of statistics. 2001; p. 1165–1188.

Benlloch-González M, Arquero O, Fournier JM, Barranco D, Benlloch M. K+ starvation inhibits water-stress-induced stomatal closure. Journal of plant physiology. 2008; 165(6):623–630.

Berg JJ, Coop G. A population genetic signal of polygenic adaptation. PLoS genetics. 2014; 10(8):e1004412.

Bolger AM, Lohse M, Usadel B. Trimmomatic: a flexible trimmer for Illumina sequence data. Bioinformatics. 2014; 30(15):2114–2120.

Bolte A, Villanueva I. Interspecific competition impacts on the morphology and distribution of fine roots in European beech (Fagus sylvatica L.) and Norway spruce (Picea abies (L.) Karst.). European Journal of Forest Research. 2006; 125:15–26.

Brang P, Pluess A, Bürgi A, Born J, Augustin S. Wald im Klimawandel. Grundlagen für Adaptationsstrategien.. 2016;.

Braun S, Hopf SE, Mainiero R. Einfluss des Klimawandels auf Wasserbeziehungen verschiedener Waldbau-marten.. 2019;.

Braun S, Hopf SE, Tresch S, Remund J, Schindler C. 37 Years of Forest Monitoring in Switzerland: Drought Effects on Fagus sylvatica. Frontiers in Forests and Global Change. 2021 oct; 4(October):1–10. https://www.frontiersin.org/articles/10.3389/ffgc.2021.765782/full, doi: 10.3389/ffgc.2021.765782.

Braun S, Remund J, Rihm B. Indikatoren zur Schätzung des Trockenheitsrisikos in Buchen-und Fichtenwäldern. Schweizerische Zeitschrift fur Forstwesen. 2015; 166(6):361–371.

Braun S, Schindler C, Rihm B. Foliar Nutrient Concentrations of European Beech in Switzerland: Relations With Nitrogen Deposition, Ozone, Climate and Soil Chemistry. Frontiers in Forests and Global Change. 2020 mar; (March):1–15. https://www.frontiersin.org/article/10.3389/ffgc.2020.00033/full, doi: 10.3389/ffgc.2020.00033.

Braun S, Tresch S, Hopf SE, Schindler C. Four Decades of Forest Development in the Context of Nitrogen Eutrophication and Climate Change. In: Springer; 2025.

Braun S, de Witte LC, Hopf SE. Auswirkungen des Trockensommers 2018 auf Flächen der Interkantonalen Walddauerbeobachtung. Schweizerische Zeitschrift fur Forstwesen. 2020; 171(5):270–280.

Breiman L. Random forests. Machine learning. 2001; 45:5–32.

Broad Institute, Picard Tools; (Accessed: 2018/02/21; version 2.17.8). http://broadinstitute.github.io/picard/.

Buras A, Rammig A, Zang CS. Quantifying impacts of the 2018 drought on European ecosystems in comparison to 2003. Biogeosciences. 2020; 17(6):1655–1672.

Cardoso AA, Visel D, Kane CN, Batz TA, García Sánchez C, Kaack L, Lamarque LJ, Wagner Y, King A, Torres-Ruiz JM, et al. Drought-induced lacuna formation in the stem causes hydraulic conductance to decline before xylem embolism in Selaginella. New Phytologist. 2020; 227(6):1804–1817.

Carter GA. Primary and secondary effects of water content on the spectral reflectance of leaves. American journal of botany. 1991; 78(7):916–924.

Cavender-Bares J, Meireles JE, Couture JJ, Kaproth MA, Kingdon CC, Singh A, Serbin SP, Center A, Zuniga E, Pilz G, et al. Associations of leaf spectra with genetic and phylogenetic variation in oaks: prospects for remote detection of biodiversity. Remote Sensing. 2016; 8(3):221.

Ceppi MG, Oukarroum A, Çiçek N, Strasser RJ, Schansker G. The IP amplitude of the fluorescence rise OJIP is sensitive to changes in the photosystem I content of leaves: a study on plants exposed to magnesium and sulfate deficiencies, drought stress and salt stress. Physiologia plantarum. 2012; 144(3):277–288.

Chen K, Zhang F, Kan J. Characterization of chlorophyll breakdown in green prickleyashes (Zanthoxylum schini-folium Zucc.) during slow drying. European Food Research and Technology. 2012; 234:1023–1031.

Chen RH, Chen YH, Huang TY. Ubiquitin-mediated regulation of autophagy. Journal of Biomedical Science. 2019; 26:1–12.

Choat B, Brodribb TJ, Brodersen CR, Duursma RA, López R, Medlyn BE. Triggers of tree mortality under drought. Nature. 2018; 558(7711):531–539.

Churchill GA, Doerge R. Empirical threshold values for quantitative trait mapping. Genetics. 1994; 138(3):963–971.

Csilléry K, Lalagüe H, Vendramin GG, González-Martínez SC, Fady B, Oddou-Muratorio S. Detecting short spatial scale local adaptation and epistatic selection in climate-related candidate genes in E uropean beech (F agus sylvatica) populations. Molecular Ecology. 2014; 23(19):4696–4708.

Curran PJ. Remote sensing of foliar chemistry. Remote sensing of environment. 1989; 30(3):271–278.

Czyż EA, Schmid B, Eppinga MB, Harpe MdL, Moradi A, Li C, Schaepman ME, Schuman MC, Inferring genetic structure of European beech from observations of spectral phenotypes. bioRxiv; 2023. https://www.biorxiv.org/content/10.1101/2023.08.16.553487v1, doi: 10.1101/2023.08.16.553487, pages: 2023.08.16.553487 Section: New Results.

Danecek P, Bonfield JK, Liddle J, Marshall J, Ohan V, Pollard MO, Whitwham A, Keane T, McCarthy SA, Davies RM, et al. Twelve years of SAMtools and BCFtools. Gigascience. 2021; 10(2):giab008.

Devlin B, Roeder K. Genomic control for association studies. Biometrics. 1999; 55(4):997–1004.

Dietrich L, Delzon S, Hoch G, Kahmen A. No role for xylem embolism or carbohydrate shortage in temperate trees during the severe 2015 drought. Journal of Ecology. 2019; 107(1):334–349.

Fitzgerald TL, Waters DL, Henry RJ. Betaine aldehyde dehydrogenase in plants. Plant biology. 2009; 11(2):119–130.

Fox EA, Wright AE, Fumagalli M, Vieira FG. ngsLD: evaluating linkage disequilibrium using genotype likelihoods. Bioinformatics. 2019; 35(19):3855–3856.

Frei ER, Gossner MM, Vitasse Y, Queloz V, Dubach V, Gessler A, Ginzler C, Hagedorn F, Meusburger K, Moor M, Samblás Vives E, Rigling A, Uitentuis I, Von Arx G, Wohlgemuth T. European beech dieback after premature leaf senescence during the 2018 drought in northern Switzerland. Plant Biology. 2022 Dec; 24(7):1132–1145. https://onlinelibrary.wiley.com/doi/10.1111/plb.13467, doi: 10.1111/plb.13467.

Futschik A, Schlotterer C. The next generation of molecular markers from massively parallel sequencing of pooled DNA samples. Genetics. 2010; 186(1):207–218.

Gárate-Escamilla H, Hampe A, Vizcaíno-Palomar N, Robson TM, Benito Garzón M. Range-wide variation in local adaptation and phenotypic plasticity of fitness-related traits in Fagus sylvatica and their implications under climate change. Global Ecology and Biogeography. 2019; 28(9):1336–1350.

Gloss AD, Vergnol A, Morton TC, Laurin PJ, Roux F, Bergelson J. Genome-wide association mapping within a local Arabidopsis thaliana population more fully reveals the genetic architecture for defensive metabolite diversity. Philosophical Transactions of the Royal Society B: Biological Sciences. 2022 May; 377(1855):20200512. https://royalsocietypublishing.org/doi/10.1098/rstb.2020.0512, doi: 10.1098/rstb.2020.0512, publisher: Royal Society.

Grandbastien MA. Activation of plant retrotransposons under stress conditions. Trends in plant science. 1998; 3(5):181–187.

Grandbastien MA. LTR retrotransposons, handy hitchhikers of plant regulation and stress response. Biochimica et Biophysica Acta (BBA)-Gene Regulatory Mechanisms. 2015; 1849(4):403–416.

Gurney A, Manoury B. Two-pore potassium channels in the cardiovascular system. European Biophysics Journal. 2009; 38:305–318.

Haworth M, Centritto M, Giovannelli A, Marino G, Proietti N, Capitani D, De Carlo A, Loreto F. Xylem morphology determines the drought response of two Arundo donax ecotypes from contrasting habitats. Gcb Bioenergy. 2017; 9(1):119–131.

Hertel D, Strecker T, Müller-Haubold H, Leuschner C. Fine root biomass and dynamics in beech forests across a precipitation gradient–is optimal resource partitioning theory applicable to water-limited mature trees? Journal of Ecology. 2013; 101(5):1183–1200.

Hong-Bo S, Li-Ye C, Ming-An S. Calcium as a versatile plant signal transducer under soil water stress. BioEssays. 2008; 30(7):634–641.

ICP Forests, Manual on methods and criteria for hamonized sampling, assessment, monitoring and analysis of the effects of air pollution on forests. Part XIV. Hamburg: Johann Heinrich von Thünen Institute; 2022.

Ilyas M, Nisar M, Khan N, Hazrat A, Khan AH, Hayat K, Fahad S, Khan A, Ullah A. Drought tolerance strategies in plants: a mechanistic approach. Journal of Plant Growth Regulation. 2021; 40:926–944.

Jabeen N, Ahmad R, Sultana R, Saleem R, Ambrat. Investigations on foliar spray of boron and manganese on oil content and concentrations of fatty acids in seeds of sunflower plant raised through saline water irrigation. Journal of plant nutrition. 2013; 36(6):1001–1011.

Jacquemoud S, Ustin S. Leaf optical properties. Cambridge: Cambridge University Press; 2019.

Jacquemoud S, Ustin SL. Leaf optical properties: A state of the art. In: 8th International Symposium of Physical Measurements & Signatures in Remote Sensing CNES Aussois France; 2001. p. 223–332.

Joseph J, Luster J, Bottero A, Buser N, Baechli L, Sever K, Gessler A. Effects of drought on nitrogen uptake and carbon dynamics in trees. Tree physiology. 2021; 41(6):927–943.

Klesse S, Wohlgemuth T, Meusburger K, Vitasse Y, von Arx G, Lévesque M, Neycken A, Braun S, Dubach V, Gessler A, et al. Long-term soil water limitation and previous tree vigor drive local variability of drought-induced crown dieback in Fagus sylvatica. Science of the Total Environment. 2022; 851:157926.

Knight H, Trewavas AJ, Knight MR. Calcium signalling in Arabidopsis thaliana responding to drought and salinity. The Plant Journal. 1997; 12(5):1067–1078.

Kofler R, Pandey RV, Schlötterer C. PoPoolation2: identifying differentiation between populations using sequencing of pooled DNA samples (Pool-Seq). Bioinformatics. 2011; 27(24):3435–3436.

Kohli A, Narciso JO, Miro B, Raorane M. Root proteases: reinforced links between nitrogen uptake and mobilization and drought tolerance. Physiologia Plantarum. 2012; 145(1):165–179.

Kokot M, Długosz M, Deorowicz S. KMC 3: counting and manipulating k-mer statistics. Bioinformatics. 2017; 33(17):2759–2761.

Korneliussen TS, Albrechtsen A, Nielsen R. ANGSD: Analysis of Next Generation Sequencing Data. BMC Bioinformatics. 2014 Nov; 15(1):356. http://www.biomedcentral.com/1471-2105/15/356/abstract, doi: 10.1186/s12859-014-0356-4.

Kramer K, Degen B, Buschbom J, Hickler T, Thuiller W, Sykes MT, de Winter W. Modelling exploration of the future of European beech (Fagus sylvatica L.) under climate change—range, abundance, genetic diversity and adaptive response. Forest Ecology and Management. 2010; 259(11):2213–2222.

Kumar J, Yumnam S, Basu T, Ghosh A, Garg G, Karthikeyan G, Sengupta S. Association of polymorphisms in 9p21 region with CAD in North Indian population: replication of SNPs identified through GWAS. Clinical genetics. 2011; 79(6):588–593.

Lalagüe H, Csilléry K, Oddou-Muratorio S, Safrana J, De Quattro C, Fady B, González-Martínez S, Vendramin G. Nucleotide diversity and linkage disequilibrium at 58 stress response and phenology candidate genes in a European beech (Fagus sylvatica L.) population from southeastern France. Tree Genetics & Genomes. 2014; 10:15–26.

Langmead B, Salzberg SL. Fast gapped-read alignment with Bowtie 2. Nature methods. 2012; 9(4):357–359.

Lazic D, Geßner C, Liepe KJ, Lesur-Kupin I, Mader M, Blanc-Jolivet C, Gömöry D, Liesebach M, González-Martínez SC, Fladung M, Degen B, Müller NA. Genomic variation of European beech reveals signals of local adaptation despite high levels of phenotypic plasticity. Nature Communications. 2024 Oct; 15(1):8553. https://www.nature.com/articles/s41467-024-52933-y, doi: 10.1038/s41467-024-52933-y, publisher: Nature Publishing Group.

Li C, Czyż EA, Schmid B, Ray R, Halitschke R, Baldwin IT, Schaepman ME, Schuman MC. Genome-wide associations of leaf spectral variation in MAGIC lines of Nicotiana attenuata. Ecology. 2026; 107(4):e70366.

Liaw A, Wiener M, et al. Classification and regression by randomForest. R news. 2002; 2(3):18–22.

Lillesaeter O. Spectral reflectance of partly transmitting leaves: Laboratory measurements and mathematical modeling. Remote Sensing of Environment. 1982 Jul; 12(3):247–254. https://doi.org/10.1016/0034-4257(82)90057-8, doi: 10.1016/0034-4257(82)90057-8.

Liu H, Stone SL. Abscisic acid increases Arabidopsis ABI5 transcription factor levels by promoting KEG E3 ligase self-ubiquitination and proteasomal degradation. The Plant Cell. 2010; 22(8):2630–2641.

Lou RN, Jacobs A, Wilder AP, Therkildsen NO. A beginner’s guide to low-coverage whole genome sequencing for population genomics. Molecular Ecology. 2021; (July):5966–5993. https://doi.org/10.1111/mec.16077, doi: 10.1111/mec.16077.

van der Maaten E, Stolz J, Thurm EA, Schröder J, Henkel A, Leinemann L, Profft I, Voth W, van der Maaten-Theunissen M. Long-term growth decline is not reflected in crown condition of European beech after a recent extreme drought. Forest Ecology and Management. 2024 Jan; 551:121516. https://www.sciencedirect.com/science/article/pii/S0378112723007508, doi: 10.1016/j.foreco.2023.121516.

Magri D. Patterns of post-glacial spread and the extent of glacial refugia of European beech (Fagus sylvatica). Journal of Biogeography. 2008; 35(3):450–463.

Mann HB, Whitney DR. On a test of whether one of two random variables is stochastically larger than the other. The annals of mathematical statistics. 1947; p. 50–60.

Mantel N. Chi-square tests with one degree of freedom; extensions of the Mantel-Haenszel procedure. Journal of the American Statistical Association. 1963; 58(303):690–700.

Mantel N, Haenszel W. Statistical aspects of the analysis of data from retrospective studies of disease. Journal of the national cancer institute. 1959; 22(4):719–748.

Marjoram P, Zubair A, Nuzhdin S. Post-GWAS: where next? More samples, more SNPs or more biology? Heredity. 2014; 112(1):79–88.

Marschner H. Marschner’s mineral nutrition of higher plants. Academic press; 2011.

McKnight PE, Najab J. Mann-Whitney U Test. The Corsini encyclopedia of psychology. 2010; p. 1–1.

Meier IC, Leuschner C. Genotypic variation and phenotypic plasticity in the drought response of fine roots of European beech. Tree physiology. 2008; 28(2):297–309.

Meireles JE, Schweiger AK, Cavender-Bares JM. spectrolab: Class and Methods for Hyperspectral Data. R pack-age version 0.0. 2. 2017;.

Meisner J, Albrechtsen A. Inferring population structure and admixture proportions in low-depth NGS data. Genetics. 2018; 210(2):719–731.

Milesi P, Kastally C, Dauphin B, Cervantes S, Bagnoli F, Budde KB, Cavers S, Fady B, Faivre-Rampant P, González-Martínez SC, Grivet D, Gugerli F, Jorge V, Lesur Kupin I, Ojeda DI, Olsson S, Opgenoorth L, Pinosio S, Plomion C, Rellstab C, et al. Resilience of genetic diversity in forest trees over the Quaternary. Nature Communications. 2024 Oct; 15(1):8538. https://www.nature.com/articles/s41467-024-52612-y, doi: 10.1038/s41467-024-52612-y, publisher: Nature Publishing Group.

Miller J, Steven M, Demetriades-Shah T. Reflection of layered bean leaves over different soil backgrounds: measured and simulated spectra. International Journal of Remote Sensing. 1992; 13(17):3273–3286.

Mishra B, Ulaszewski B, Meger J, Aury JM, Bodénès C, Lesur-Kupin I, Pfenninger M, Da Silva C, Gupta DK, Gui-choux E, et al. A Chromosome-level genome assembly of the European Beech (Fagus sylvatica) reveals anomalies for organelle DNA integration, repeat Content and distribution of SNPs. Frontiers in genetics. 2022; 12:691058.

Mueller N, Lewis A, Roberts D, Ring S, Melrose R, Sixsmith J, Lymburner L, McIntyre A, Tan P, Curnow S, et al. Water observations from space: Mapping surface water from 25 years of Landsat imagery across Australia. Remote Sensing of Environment. 2016; 174:341–352.

Mukherjee R, Das A, Chakrabarti S, Chakrabarti O. Calcium dependent regulation of protein ubiquitination– Interplay between E3 ligases and calcium binding proteins. Biochimica et Biophysica Acta (BBA)-Molecular Cell Research. 2017; 1864(7):1227–1235.

Muttenz PH, Liebendörfer H, Meier H. Heimatkunde Muttenz - Der Hardwald. In: Heimatkunde Muttenz 2009 - zu Beginn des neuen Jahrtausends Muttenz: Einwohnergemeinde Muttenz 2009, Verlag des Kantons Basel-Landschaft; 2009.https://www.heimatkunde-muttenz.ch/natur-und-landschaft/einzelraeume/hardwald/der-hardwald, https://www.muttenz.ch/onlineshop/27062.

Nabiyouni M, Brückner T, Zhou H, Gbureck U, Bhaduri SB. Magnesium-based bioceramics in orthopedic applications. Acta biomaterialia. 2018; 66:23–43.

Neycken A, Scheggia M, Bigler C, Lévesque M. Long-term growth decline precedes sudden crown dieback of European beech. Agricultural and Forest Meteorology. 2022; 324:109103.

Nussbaumer A, Meusburger K, Schmitt M, Waldner P, Gehrig R, Haeni M, Rigling A, Brunner I, Thimonier A. Extreme summer heat and drought lead to early fruit abortion in European beech. Scientific Reports. 2020; 10(1):5334.

Paliyath G, Murr DP, Handa AK, Lurie S. Postharvest biology and technology of fruits, vegetables, and flowers. John Wiley & Sons; 2009.

Peñuelas J, Gamon J, Fredeen A, Merino J, Field C. Reflectance indices associated with physiological changes in nitrogen-and water-limited sunflower leaves. Remote sensing of Environment. 1994; 48(2):135–146.

Petibon F, Czyż EA, Ghielmetti G, Hueni A, Kneubühler M, Schaepman ME, Schuman MC. Uncertainties in measurements of leaf optical properties are small compared to the biological variation within and between individuals of European beech. Remote Sensing of Environment. 2021 Oct; 264:112601. https://www.sciencedirect.com/science/article/pii/S0034425721003217, doi: 10.1016/j.rse.2021.112601.

Pfenninger M, Reuss F, KIebler A, Schönnenbeck P, Caliendo C, Gerber S, Cocchiararo B, Reuter S, Blüthgen N, Mody K, et al. Correction: Genomic basis for drought resistance in European beech forests threatened by climate change. ELife. 2024; 13:e102872.

Pfenninger M, Reuss F, Kiebler A, Schönnenbeck P, Caliendo C, Gerber S, Cocchiararo B, Reuter S, Blüthgen N, Mody K, Mishra B, Bálint M, Thines M, Feldmeyer B. Genomic basis for drought resistance in European beech forests threatened by climate change. eLife. 2021; 10:e65532. https://elifesciences.org/articles/65532, doi: 10.7554/eLife.65532.

Pluess AR, Frank A, Heiri C, Lalagüe H, Vendramin GG, Oddou-Muratorio S. Genome–environment association study suggests local adaptation to climate at the regional scale in Fagus sylvatica. New Phytologist. 2016; 210(2):589–601.

Poplin R, Ruano-Rubio V, DePristo MA, Fennell TJ, Carneiro MO, Van der Auwera GA, Kling DE, Gauthier LD, Levy-Moonshine A, Roazen D, et al. Scaling accurate genetic variant discovery to tens of thousands of samples. BioRxiv. 2017; p. 201178.

Purcell S, Neale B, Todd-Brown K, Thomas L, Ferreira MA, Bender D, Maller J, Sklar P, De Bakker PI, Daly MJ, et al. PLINK: a tool set for whole-genome association and population-based linkage analyses. The American journal of human genetics. 2007; 81(3):559–575.

R Core Team. R: A Language and Environment for Statistical Computing. R Foundation for Statistical Computing, Vienna, Austria; 2023, https://www.R-project.org/.

Rellstab C, Zoller S, Tedder A, Gugerli F, Fischer MC. Validation of SNP Allele Frequencies Determined by Pooled Next-Generation Sequencing in Natural Populations of a Non-Model Plant Species. PLOS ONE. 2013 Nov; 8(11):e80422. https://journals.plos.org/plosone/article?id=10.1371/journal.pone.0080422, doi: 10.1371/journal.pone.0080422, publisher: Public Library of Science.

Rieder JS, Žmegač A, Link RM, Köthe K, Ullmann T, Seidel D, Fäth J, Zang C, Schuldt B. Tree size, neighbour-hood composition and structure affect individual tree vitality of European beech following extreme drought. Forest Ecology and Management. 2026 Jan; 599:123293. https://www.sciencedirect.com/science/article/pii/S0378112725008011, doi: 10.1016/j.foreco.2025.123293.

Ruehr NK, Offermann CA, Gessler A, Winkler JB, Ferrio JP, Buchmann N, Barnard RL. Drought effects on allocation of recent carbon: from beech leaves to soil CO2 efflux. New Phytologist. 2009; 184(4):950–961.

Sanchez Herrero JF, Pluvinet R, Luna de Haro A, Sumoy L. Paired-end small RNA sequencing reveals a possible overestimation in the isomiR sequence repertoire previously reported from conventional single read data analysis. BMC bioinformatics. 2021; 22(1):1–16.

Santure AW, Garant D. Wild GWAS—association mapping in natural populations. Molecular ecology resources. 2018; 18(4):729–738.

Sardans J, Peñuelas J. Potassium control of plant functions: Ecological and agricultural implications. Plants. 2021; 10(2):419.

Saunders A, Drew DM. Measurements done on excised stems indicate that hydraulic recovery can be an important strategy used by Eucalyptus hybrids in response to drought. Trees. 2021; p. 1–13.

Schlötterer C, Tobler R, Kofler R, Nolte V. Sequencing pools of individuals—mining genome-wide polymorphism data without big funding. Nature Reviews Genetics. 2014; 15(11):749–763.

Schmid-Siegert E, Sarkar N, Iseli C, Calderon S, Gouhier-Darimont C, Chrast J, Cattaneo P, Schütz F, Farinelli L, Pagni M, et al. Low number of fixed somatic mutations in a long-lived oak tree. Nature Plants. 2017; 3(12):926–929.

Schuldt B, Buras A, Arend M, Vitasse Y, Beierkuhnlein C, Damm A, Gharun M, Grams TE, Hauck M, Hajek P, et al. A first assessment of the impact of the extreme 2018 summer drought on Central European forests. Basic and Applied Ecology. 2020; 45:86–103.

Sehar S, Adil MF, Zeeshan M, Holford P, Cao F, Wu F, Wang Y. Mechanistic insights into potassium-conferred drought stress tolerance in cultivated and tibetan wild barley: Differential osmoregulation, nutrient retention, secondary metabolism and antioxidative defense capacity. International Journal of Molecular Sciences. 2021; 22(23):13100.

Senf C, Buras A, Zang C, Rammig A, Seidl R. A consistent link between drought and forest diebacks across Europe. In: EGU General Assembly Conference Abstracts; 2020. p. 5679.

Shu K, Yang W. E3 ubiquitin ligases: ubiquitous actors in plant development and abiotic stress responses. Plant and Cell Physiology. 2017; 58(9):1461–1476.

Stefanini C, Csilléry K, Ulaszewski B, Burczyk J, Schaepman ME, Schuman MC. A novel synthesis of two decades of microsatellite studies on European beech reveals decreasing genetic diversity from glacial refugia. Tree Genetics & Genomes. 2023; 19(1):3. https://doi.org/10.1007/s11295-022-01577-4, doi: 10.1007/s11295-022-01577-4.

Storey JD, Tibshirani R. Statistical significance for genomewide studies. Proceedings of the National Academy of Sciences. 2003; 100(16):9440–9445.

Tränkner M, Tavakol E, Jákli B. Functioning of potassium and magnesium in photosynthesis, photosynthate translocation and photoprotection. Physiologia plantarum. 2018; 163(3):414–431.

Tyree MT, Cochard H, Cruiziat P, Sinclair B, Ameglio T. Drought-induced leaf shedding in walnut: evidence for vulnerability segmentation. Plant, Cell & Environment. 1993; 16(7):879–882.

Tyree MT, Sperry JS. Do woody plants operate near the point of catastrophic xylem dysfunction caused by dynamic water stress? Answers from a model. Plant physiology. 1988; 88(3):574–580.

Vitasse Y, Bottero A, Cailleret M, Bigler C, Fonti P, Gessler A, Lévesque M, Rohner B, Weber P, Rigling A, et al. Contrasting resistance and resilience to extreme drought and late spring frost in five major European tree species. Global Change Biology. 2019; 25(11):3781–3792.

Voichek Y, Weigel D. Identifying genetic variants underlying phenotypic variation in plants without complete genomes. Nature genetics. 2020; 52(5):534–540.

Voorman A, Lumley T, McKnight B, Rice K. Behavior of QQ-plots and genomic control in studies of gene-environment interaction. PloS one. 2011; 6(5):e19416.

Wang F, Israel D, Ramírez-Valiente JA, Sánchez-Gómez D, Aranda I, Aphalo PJ, Robson TM. Seedlings from marginal and core populations of European beech (Fagus sylvatica L.) respond differently to imposed drought and shade. Trees - Structure and Function. 2021; 35(1):53–67. https://doi.org/10.1007/s00468-020-02011-9, doi: 10.1007/s00468-020-02011-9, publisher: Springer Berlin Heidelberg ISBN: 0046802002.

Wang X, Komatsu S. Proteomic approaches to uncover the flooding and drought stress response mechanisms in soybean. Journal of Proteomics. 2018; 172:201–215.

Wang X, Guo L, Wang G, Li M. PLD: phospholipase Ds in plant signaling. Springer; 2014.

Wang Z, Chlus A, Geygan R, Ye Z, Zheng T, Singh A, Couture JJ, Cavender-Bares J, Kruger EL, Townsend PA. Foliar functional traits from imaging spectroscopy across biomes in eastern North America. New Phytologist. 2020 Oct; 228(2):494–511. https://onlinelibrary.wiley.com/doi/10.1111/nph.16711, doi: 10.1111/nph.16711.

Waraich EA, Ahmad R, Ashraf M. Role of mineral nutrition in alleviation of drought stress in plants. Australian Journal of Crop Science. 2011; 5(6):764–777.

Weretilnyk EA, Hanson AD. Molecular cloning of a plant betaine-aldehyde dehydrogenase, an enzyme implicated in adaptation to salinity and drought. Proceedings of the National Academy of Sciences. 1990; 87(7):2745–2749.

Wickham H. ggplot2: Elegant Graphics for Data Analysis. Springer-Verlag New York; 2016. https://ggplot2.tidyverse.org.

Wilcoxon F. Individual comparisons by ranking methods. In: Breakthroughs in statistics: Methodology and distribution Springer; 1992.p. 196–202.

Yachandra VK, Sauer K, Klein MP. Manganese cluster in photosynthesis: where plants oxidize water to dioxygen. Chemical Reviews. 1996; 96(7):2927–2950.

Zang U, Goisser M, Häberle KH, Matyssek R, Matzner E, Borken W. Effects of drought stress on photosynthesis, rhizosphere respiration, and fine-root characteristics of beech saplings: A rhizotron field study. Journal of plant nutrition and soil science. 2014; 177(2):168–177.

Zhao J, Zhao L, Zhang M, Zafar SA, Fang J, Li M, Zhang W, Li X. Arabidopsis E3 ubiquitin ligases PUB22 and PUB23 negatively regulate drought tolerance by targeting ABA receptor PYL9 for degradation. International journal of molecular sciences. 2017; 18(9):1841.

Zou JJ, Wei FJ, Wang C, Wu JJ, Ratnasekera D, Liu WX, Wu WH. Arabidopsis calcium-dependent protein kinase CPK10 functions in abscisic acid-and Ca2+-mediated stomatal regulation in response to drought stress. Plant physiology. 2010; 154(3):1232–1243.

